# Proteolytic degradation of atrial sarcomere proteins underlies contractile defects in atrial fibrillation

**DOI:** 10.1101/2023.11.05.565691

**Authors:** Hannah E Cizauskas, Hope V Burnham, Azaria Panni, Alexandra Pena, Alejandro Alvarez-Arce, M Therese Davis, Kelly N Araujo, Christine Delligatti, Seby Edassery, Jonathan A Kirk, Rishi Arora, David Y Barefield

**Author notes:** To whom correspondence should be addressed David Y. Barefield, PhD, Department of Cell and Molecular Physiology, Loyola University Chicago, 2160 S. 1st Ave, Maywood, IL 60153.

## Abstract

**Aims:** Atrial fibrillation (AFib) is the most common cardiac rhythm disturbance. Treatment of AFib involves restoration of the atrial electrical rhythm. Following rhythm restoration, a period of depressed mechanical function known as atrial stunning occurs that involves decreased blood flow velocity and reduced atrial contractility. This suggests that defects in contractility occur in AFib and are revealed upon restoration of rhythm. The aim of this project is to define the contractile remodeling that occurs in AFib

**Methods and Results:** To assess contractile function, we used a canine atrial tachypacing model of induced AFib. Mass spectrometry analysis showed dysregulation of contractile proteins in samples from AFib compared to sinus rhythm atria. Atrial cardiomyocytes showed reduced force of contraction in skinned single cardiomyocyte calcium-force studies. There were no significant differences in myosin heavy chain isoform expression. Resting tension is decreased in the AFib samples correlating with reduced full-length titin in the sarcomere. We measured degradation of other myofilament proteins including cMyBP-C, actinin, and cTnI, showing significant degradation in the AFib samples compared to sinus rhythm atria. Many of the protein degradation products appeared as discrete cleavage products that are generated by calpain proteolysis. We assessed calpain activity and found it to be significantly increased. Skinned cardiomyocytes from AFib atria showed decreased troponin I phosphorylation, consistent with the increased calcium sensitivity that was found within these cardiomyocytes.

**Conclusions:** With these results it can be concluded that AFib causes alterations in contraction that can be explained by both molecular changes occurring in myofilament proteins and overall myofilament protein degradation. These results provide an understanding of the contractile remodeling that occurs in AFib and provides insight into the molecular explanation for atrial stunning and the increased risk of atrial thrombus and stroke in AFib.

## (iii) INTRODUCTION

Atrial fibrillation (AFib) is the most common cardiac rhythm disturbance with an annual occurrence of 1 – 2%, and this incidence rate increases with age (1). About 2.3 million Americans are affected by AFib and this number is expected to increase to 6 million by 2050 (1, 2). AFib is known for both electrophysiological and fibrotic remodeling, both of which cause defects in action potential conduction (3, 4). Increased fibrosis of the atria leads to a slowing of the electrical signal propagation, creating reentrant pathways and enhancing the ability for early after depolarizations to occur (5-7).

AFib is a complex disease with many known causes and is a comorbidity in many cardiovascular and non-cardiovascular diseases. Although we have gained an advanced understanding of the electrophysiology and morphological changes that cause AFib, little is known about the relationship between AFib and the contractile apparatus and contractile function. It has been established that primary contractile dysfunction can cause and be exacerbated by AFib (8, 9). Furthermore, atrial contractile dysfunction, and the resulting morphological and hemodynamic changes within the atria, is central to the increased risk of thrombus formation in the left atrial appendage, and the resulting embolic stroke risk associated with AF.

Restoration of sinus rhythm from AFib via cardioversion causes a period of mechanical dysfunction known as atrial stunning (10). Atrial stunning entails minimal contractility, indicated by decreased atrial strain during active contraction, that reduces mitral blood flow velocity and atrial contribution to ventricular filling (10-12). This suggests that there are underlying changes within the contractile apparatus that occur during AFib. The resolution of atrial stunning can occur within minutes to several weeks, with the duration of stunning correlating to the duration of AFib and the presence of comorbid heart failure (15, (12, 13)). The mechanism for stunning is not well understood, but the variability in resolution suggests the mechanisms involved require significant time to recover. We hypothesized that the aberrant signaling and cellular stress that occurs in atrial cardiomyocytes causes dysregulation of contractile proteins that may contribute to both atrial stunning and the risk for thrombus development and stroke in AFib. To study the impact of AFib on contractile protein structure and function, we utilized a dog atrial tachypacing model of AFib.

## (iv) MATERIALS AND METHODS

### Animal Model

This experimental study used 12 purpose-bred hounds. Canines were assigned as either control groups in sinus rhythm (SR) (n = 6) or were subjected to rapid atrial pacing (RAP) (n = 6) as previously described to induce AFib (14). All canines were maintained in accordance with the Guide for the Care and Use of Laboratory Animals in an AAALAC-accredited facility at Northwestern University. All procedures were approved by the IACUC of Northwestern University. Through fluoroscopic guidance, a Medtronic transvenous pacing lead was screwed into the right atrial appendage and connected to an implanted, programmable generator. One week later, RAP was initiated after X-ray confirmation of lead placement (Fig. 1. A). Animals were paced at 600 pulses/min, 0.5 mV, and pulse width 0.4 ms for 16 hours a day until persistent AFib was induced. After pacemaker implantation, all RAP dogs underwent frequent pacemaker interrogations to determine the duration of induced AFib episodes for 4 weeks, typically entering sustained AFib in week 3 (Fig. 1.B). Baseline and terminal apical four chamber echocardiographs were performed on four RAP canines and control measurements from a previous study were used to determine if the RAP canines also developed left ventricular failure (15). There were no significant differences between the baseline and terminal LA volume, LV ejection fraction, septal E/e’ ratio, or Lateral E/e’ ratio, as determined via paired T-Tests. However, the low LV ejection fraction in these canines indicates that they are experiencing systolic dysfunction (Table 1). Atrial tissue was collected after a terminal electrophysiology study at the end of week four (Fig. 1.C).

**Table 1.** Rapid atrial pacing does not induce ventricular failure in the canines used for this study. Baseline and Terminal Echocardiographic measurements from 4 of the canines that underwent rapid atrial pacing (RAP). LA volume, Ejection fraction, E/e’ Septal, and E/e’ lateral values were not significantly different from the baseline and post-RAP terminal echocardiographs. Average values for these parameters from control canines that did not undergo RAP were taken from a previously published data set from the same research group (15).

|  | LA Volume | Ejection Fraction | E/e' Septal | E/e' Lateral |
| --- | --- | --- | --- | --- |
| Baseline | 19.75 ± 2.5 | 27.5 ± 6.95 | 8.735 ± 1.52 | 6.76 ± 0.4 |
| Terminal | 22.5 ± 3.57 | 18 ± 7.04 | 7.52 ± 0.54 | 5.69 ± 0.92 |
| P-Value | 0.55 | 0.37 | 0.48 | 0.33 |

**Fig. 1.**
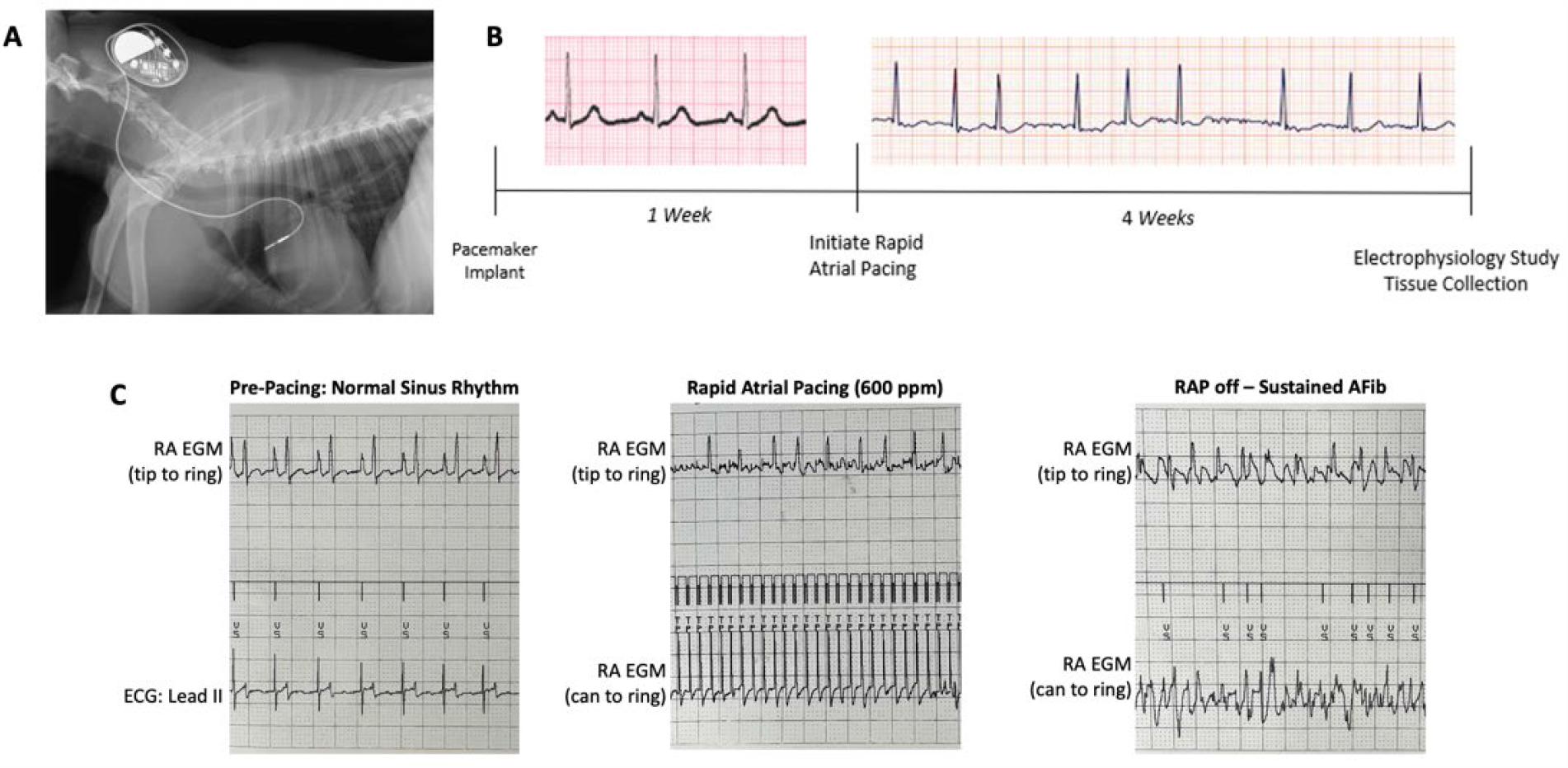
Atrial tachypacing in a canine through the right atrial appendage causes atrial fibrillation. A) X-ray confirmation of transvenous lead in the right atrial appendage. B) Timeline of atrial tachypacing. C) Sample intracardiac electrogram recordings from a RAP dog: prior to pacing (left), during pacing (center), and after sustained AFib (right).

### Mass Spectrometry and Data Analysis

Detailed methods for the mass spectrometry experiment can be found in the supplemental methods. Raw data were analyzed using Proteome Discoverer 2.5 (Thermo Fisher) using Sequest HT search engines. The data were searched against the Dog entries in the Uniprot protein sequence database (Canis lupus familiaris Proteome ID UP000002254). The search parameters included precursor mass tolerance 10 ppm and 0.6 Da for fragments, allowing 2 missed trypsin cleavages, oxidation (Met) and acetylation (protein N-term) as variable modifications, and carbamidomethylation (Cys) as a static modification. Percolator PSM validation was used with the following parameters: strict false discover rate of 0.01, relaxed false discovery rate of 0.05, maximum ΔCn of 0.05, and validation based on q-value. Precursor Ions Quantifier settings were Peptides to Use: Unique + Razor; Consider Protein Groups for Peptide Uniqueness set as True; Precursor Abundance Based On: Intensity; Normalization based on Total Peptide Amount; Scaling Mode set as none, Protein Abundance Calculation based on Top 10 Average intensity, low abundance peptides were removed by filtering out proteins with less than 3 PSMs, and T-Tests (background based) were used for class comparison. Differentially expressed proteins were selected based on p-value < 0.05 and log2 fold change > 1.0.

Volcano plots were made using ggVolcanoR (https://ggvolcanor.erc.monash.edu) by inputting the differentially expressed genes along with the logFC and p-value for each gene. The cutoff values for the volcano plot were 0.05 for the p-value and 0.58 for the logFC value. Gene ontology was analyzed using the DAVID annotation tool (DAVID: Functional Annotation Tools (ncifcrf.gov)). In this annotation tool a list of all differentially expressed genes was input and significantly different gene ontology pathways were obtained and analyzed using heat maps via HeatMapper program (http://www.heatmapper.ca/expression). In the HeatMapper program, a list of the genes involved in the gene ontology pathway along with their expressions for each sample was input and then plotted using Pearson complete linkage.

### Skinned Myocyte Permeabilization, Force Calcium and Resting Tension

Canine LA was prepared as previously. Briefly, canine LA myocytes were homogenized in Isolation solution (Table X, pH 7.02), with 0.3% Triton and protease and phosphatase inhibitors (Thermo Scientific). Homogenates incubated on ice (“skinned”) on ice for 20 minutes. After this, homogenate was centrifuged at 120xg, supernatant discarded, and pellet re-suspended and centrifuged twice in Isolation solution (without triton) to remove detergent. Individual cells were attached to micro-pins via UV-curing glue (glues NOA61, NOA68, ThorLabs), connected to a force transducer (Aurora Scientific) and length controller (Aurora Scientific). Sarcomere length was set to 2.1 μm and monitored via a video camera and high-speed video sarcomere length software (Aurora Scientific). Experiments were conducted at room temperature.

Active force was measured as previously. Cells chosen for active force were moved to relaxing solution (Table X, pH 7.02) with protease and phosphatase inhibitors. To measure force, myocytes were exposed to increasing free calcium concentrations, from near-zero to 46.8μM, obtained by mixing activating solution (Table X, pH 7.02) and relaxing solution in various ratios. Activation solution also contained protease inhibitors. Measurements were taken using Aurora Scientific Model 600A Digital Controller and Scope. In separate preparations, cells chosen for resting tension were maintained in Isolation solution. Sarcomere length was increased from 1.8 to 2.4 μm with length controlling pin and force was measured every 0.1μm of length. Cells were fit to exponential curves. Generally, approximately the same number of cells were used per animal.

The data was fit to a modified hill equation and significance was determined by student’s T-test analysis (force-calcium) or two-way ANOVA (resting tension) in PRISM version 9.

### Protein isolation and gels for Titin and Myosin

10 – 15 mg of tissue from all 12 canine atria samples (n = 6 SR, 6 AFib) was used for protein isolation. The tissue was crushed and dissolved in equal volumes of 8M urea solution and 50% glycerol solution. Protein concentration was then determined using the RC/DC protein quantification assay (Biorad, 5000122). To resolve titin, the isolated protein was run on a 16 cm by 18 cm 1% agarose gel. Myosin isoforms were resolved on a 16 cm by 18 cm 6.25% Acrylamide SDS-PAGE gel. In depth descriptions of the methods for protein isolation, titin gels, and myosin gels can be found in the supplemental methods.

All titin and myosin gels were analyzed using densitometry in ImageJ version 2.9. Significance between the SR and AFib samples was investigated using T-Tests in GraphPad PRISM v. 9.

### Myofilament fractionation and immunoblots

Myofilament fractionation and immunoblots were performed as described in the supplemental methods. Protein expression was analyzed via densitometry in ImageJ version 2.9 and significance was determined using T-Tests in GraphPad PRISM v. 9.

### Calpain Activity

Calpain activity was measured using the Abcam calpain activity assay kit, per manufacturer instructions (Abcam, ab65308). Briefly, 10 mg of tissue from all 12 samples (n = 6 SR, 6 AFib) was homogenized in the extraction buffer provided in the kit. 15 μg of protein was then mixed with 5uL of calpain substrate and incubated in the dark for 30 minutes. Fluorescence was measured at ex/em 400/505 nm to detect the relative amount of the calpain substrate that was degraded, providing a quantification of relative calpain activity. Each sample was performed in triplicate. Positive and negative controls were performed using active calpain and calpain inhibitor provided in the kit. The samples were normalized to the negative control and a T-Test was used to compare the SR and AFib samples in GraphPad PRISM v. 9.

## (v) RESULTS

### Atrial Fibrillation shows alterations in contractility genes and protein pathways

Analysis of the canine mass spec data revealed dysregulation of numerous proteins in AFib. 3% of the 3,898 proteins identified were significantly downregulated in the AFib samples and 8.8% were significantly upregulated (Tables S1 and S2). To investigate the molecular pathways that are altered in AFib, mass spectrometry data was input into the DAVID Gene Ontology (GO) Program. This revealed dysregulated GO pathways including calcium signaling, apoptosis, fibrosis, and metabolism (Fig. S1). As these pathways are commonly dysregulated in human AFib. Dysregulation of apoptosis and fibrosis are commonly not seen in smaller animal models of AFib and when seen, are reversed when pacing is stopped, including mice, indicating that this model of AFib better recapitulates the human disease. Two of the significantly altered GO pathways included cardiac muscle contraction and hypertrophic cardiomyopathy (Fig. 2A).

**Fig. 2.**
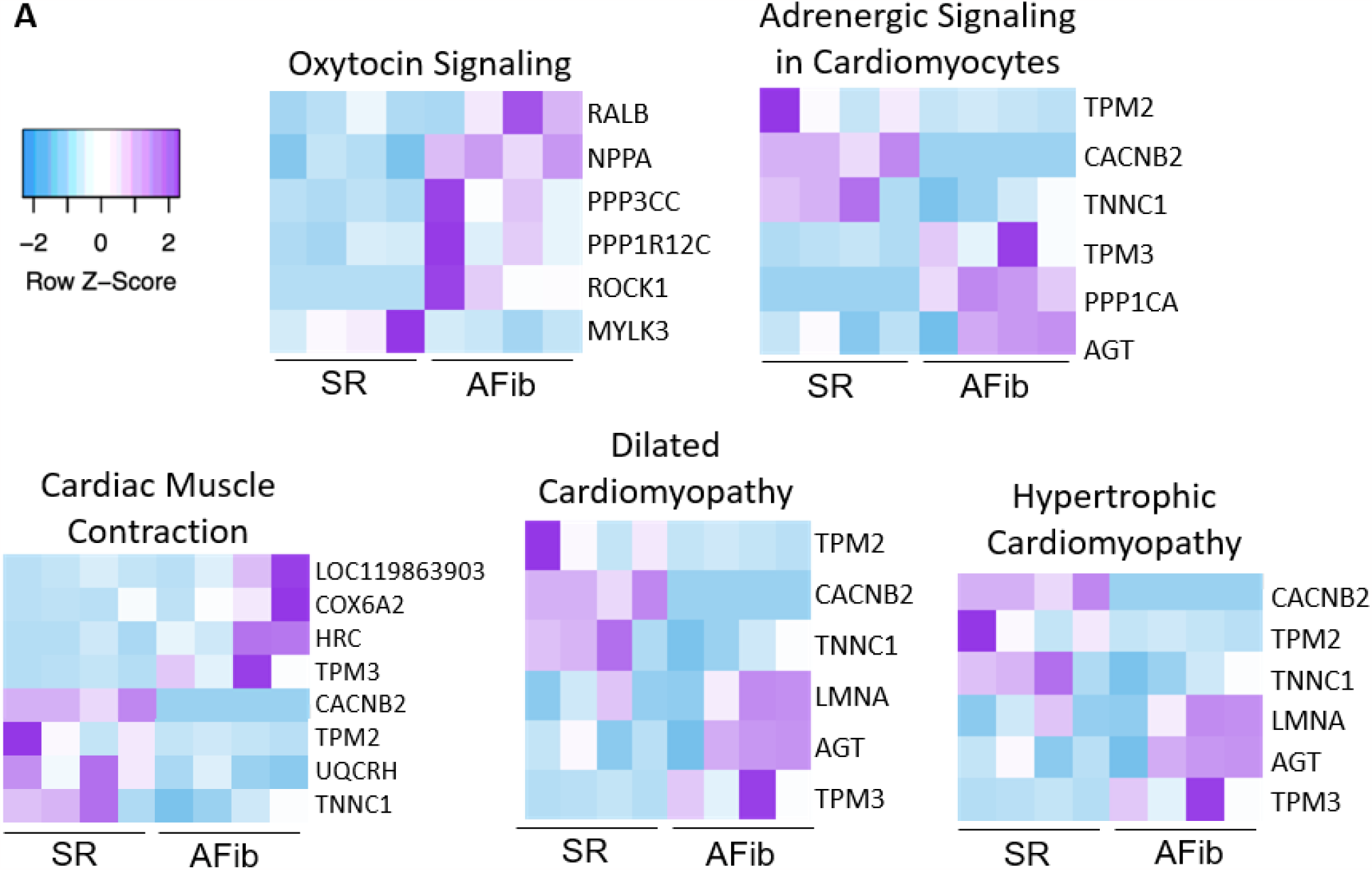
Mass Spectrometry Data shows alterations in cardiac contractility genes and protein pathways. B) Heat maps for contractility related Gene Ontology Pathways in AFib which include dilated cardiomyopathy, hypertrophic cardiomyopathy, cardiac muscle contraction, adrenergic signaling in cardiomyocytes, and oxytocin signaling.

Further investigation of contractile protein dysregulation in the AFib samples revealed alterations in proteins involved in adrenergic signaling in cardiomyocytes, oxytocin signaling, and dilated cardiomyopathy (Fig. 2B). Significantly downregulated proteins involved in contractility within the AFib samples include *MLIP, MYH7B, MYLK3, S100A1, TNNC1, TNNI1*, and *TPM2* (Table S1). Contractile proteins upregulated in the AFib samples include *ANXA6, CFL1, MPRIP, MYO1F, MYOM1, PPP1CA, PPP1R12C, PPP3CC, PRKAG3*, and *TPM3* (Table S2). Changes in expression of these proteins implicate a role for direct alterations in contractile function along with alterations in post-translational modifications of myofilament proteins that can alter contractile function.

### Atrial Fibrillation results in reduced force of contraction

Knowing that contractile proteins are dysregulated in our samples, we hypothesized that biophysical parameters of contraction will be altered in AFib. We first conducted force-calcium experiments which show a reduction in the maximum force of contraction of the AFib atrial cardiomyocytes (Fig. 3A, B). A known contribution to maximum force of contraction is MHC isoform expression. The α-MHC isoform allows for faster contraction with reduced force of contraction whereas β-MHC produces slower and more energetically efficient contractions due to their different ATPase activities (28). We hypothesized that the AFib samples would have a decrease in the percent β-MHC expression. To identify changes in MHC isoform expression, samples from both groups were run in myosin gels, followed by staining with Sypro Ruby (Fig. 3C). Isoform expression analysis using ImageJ and Prism showed no significant differences in the percent β-MHC between the two groups (Fig. 3D). To determine whether alterations in MHC head confirmation contribute to the changes in the force development in our AFib group, we performed a MantATP assay. This revealed changes in the fast and slow phase of fluorescent ATP decay and a significant increase in the percent of myosin heads in the super relaxed (SRX) conformation (Fig. S2). Increased SRX MHC head conformation may provide a potential explanation for the decrease in contractile force that is occurring in the AFib samples.

**Fig. 3.**
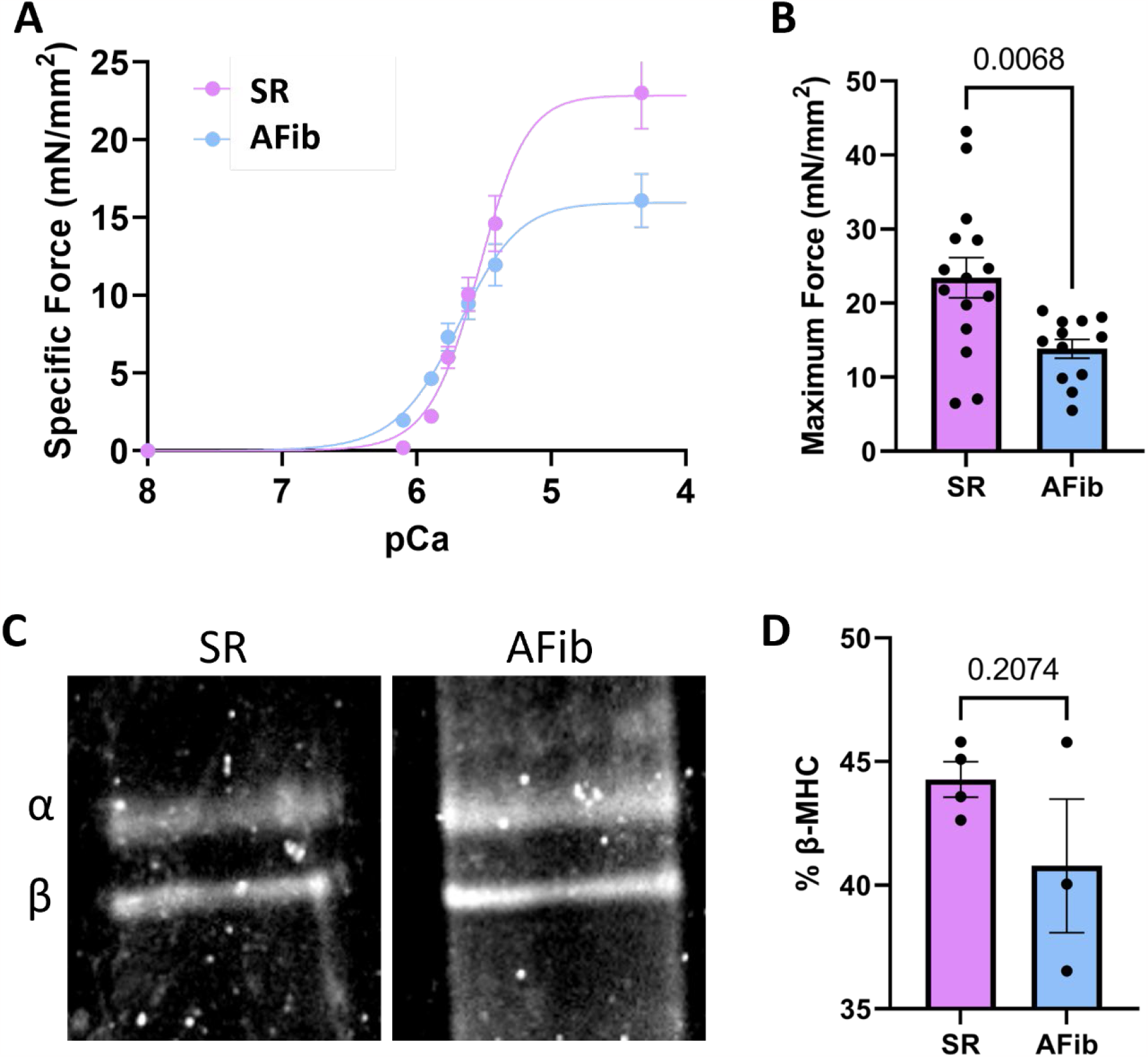
Atrial Fibrillation causes reduced contractile force. A) Skinned myocyte force-calcium relationship between SR and AFib hearts shows a reduction in maximum force. B) Summary data for myocyte max force. C) Representative images of SR (left) and AFib (right) myosin gel. D) Analysis of myosin gel shows no difference in isoform expression.

### Atrial Fibrillation resulted in reduced resting tension of atrial cardiomyocytes

We next hypothesized that another biophysical parameter of contraction, resting tension, may be altered in AFib. To investigate this, we analyzed resting tension of the atrial cardiomyocytes. Resting tension experiments revealed a reduction in the resting tension of the AFib atrial cardiomyocytes compared to the SR atrial cardiomyocytes (Fig. 4A). Because titin is one of the main proteins involved in resting tension of cardiomyocytes (21), alterations in titin can reduce resting tension. First, titin isoform expression was evaluated. We hypothesized that the AFib samples would have an increase in the N2BA/N2B ratio as N2BA is longer and produces less resting tension than N2B titin (22). There was no significant change in the N2BA/N2B expression ratio between the groups (Fig. 4B, C). We then hypothesized that there would be an increase in phosphorylation of titin in the AFib samples because phosphorylation of specific titin domains reduces resting tension (23). However, there were no significant changes in phosphorylation levels between the two groups (Fig. S3B).

**Fig. 4.**
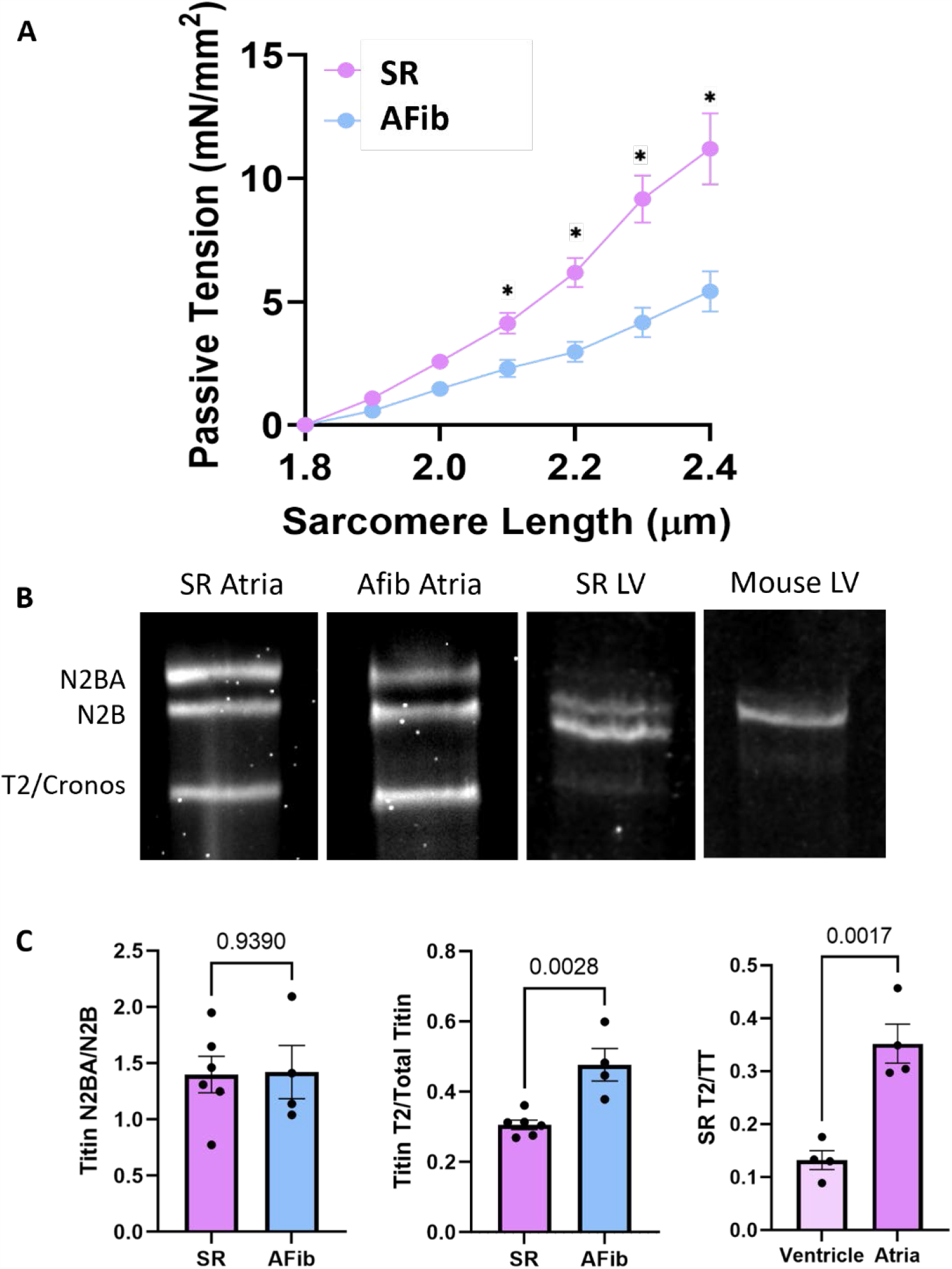
Atrial Fibrillation shows reduction in resting tension and increased titin degradation. A) Skinned atrial cardiomyocyte resting force sarcomere length and tension shows decreased resting tension in the AFib group. B) Representative images of SR atria (left), AFib atria (second), SR left ventricle (third), and mouse left ventricle (right) titin gel. C) Analysis of titin gel shows no difference in isoform expression between SR and AF groups however, there is an increase in the expression of the short titin protein (T2/Cronos) in the AFib samples. Analysis of the expression of the T2/Cronos fragment of titin in the SR ventricles and atria indicates that healthy atria express the T2/Cronos isoform at a higher ratio than the ventricles.

Analysis of the short T2/Cronos fragment of titin was normalized to total titin, which showed a significant increase in T2/Cronos levels in the AFib samples (Fig. 4C). To investigate if higher amounts of T2/Cronos occur in the atria compared to ventricle, titin extraction was performed on the SR ventricle and atria and showed that T2/Cronos is significantly more abundant in the atria than in the ventricles. To control for the possibility that the T2/Cronos fragment was generated through the titin extraction protocol, we also isolated titin from wild-type mouse ventricles, because the expression of T2/Cronos is well established in the mouse left ventricle, at the same time and showed low levels of T2/Cronos consistent with other published data in ventricular tissue (18). This indicates that the high level of T2/Cronos within the canine atria samples were not caused by the extraction, but rather an intrinsic property of the canine atria.

### Atrial Fibrillation causes increased myofilament protein degradation and increased calpain activity

Titin is known to be one of many targets for calpain degradation (24-27). Because we observed less full-length titin in the AFib samples, myofilament fractionation was performed, and myofilament gels were run to analyze protein expression and degradation products that are maintained in the sarcomere (Fig. 5A). Analysis of the myofilament gels shows distinct protein cleavage bands within the AFib samples that are absent in the SR samples. It also shows reduced protein content within the myofilaments in the AFib samples (Fig. 5B). As Calpain also targets troponin I (TnI), MHC, actinin, and myosin binding protein C (MyBPC) (24-27), these proteins were investigated for protein degradation via immunoblotting. Actinin and cMyBP-C (Fig. 5B) and TnI (Fig. 6C) showed specific degradation patterns in the AFib samples that are almost completely absent in the SR samples. Next, to determine if calpain overexpression would be altering the degradation, expression of calpain isoforms was measured via the mass spectrometry data. Mass spectrometry revealed expression of calpain 1, calpain 2, and calpain 5. Expression of calpain one, two, and five was not significantly different between the SR and AFib samples (Fig. 5C). However, calpains are activated by intracellular calcium levels. Since calcium cycling is significantly altered in AFib, leading to increase in calcium sparks and triggered waves – we next assessed calpain activity in AFib samples. Calpain activity is significantly increased in the AFib samples compared to the SR samples (Fig. 5D). This suggests that the myofilament degradation that was observed may be due to increased calpain activity.

**Fig. 5.**
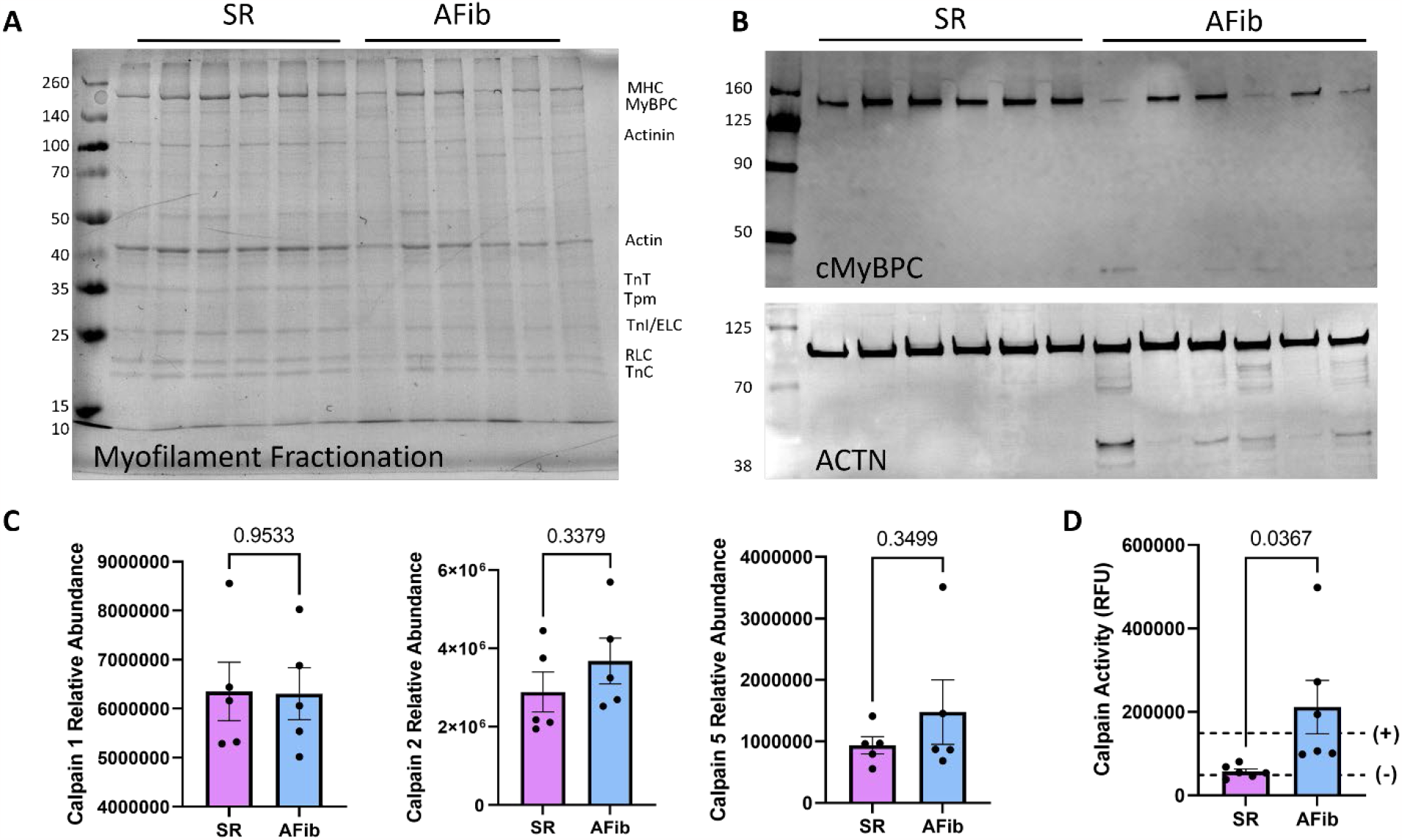
Atrial fibrillation induces calpain activity and myofilament protein degradation. A) Myofilament fractionation gel shows apparent degradation bands in the AFib samples that are absent in the AFib samples. B) Immunoblots of actinin and cMyBP-C expression show degradation patterns in the AFib samples that are almost completely absent in the SR samples. C) Expression of calpain 1, calpain 2, and calpain 5 are not statistically different in SR and AFib. D) Calpain activity is significantly increased in AFib model.

**Fig. 6.**
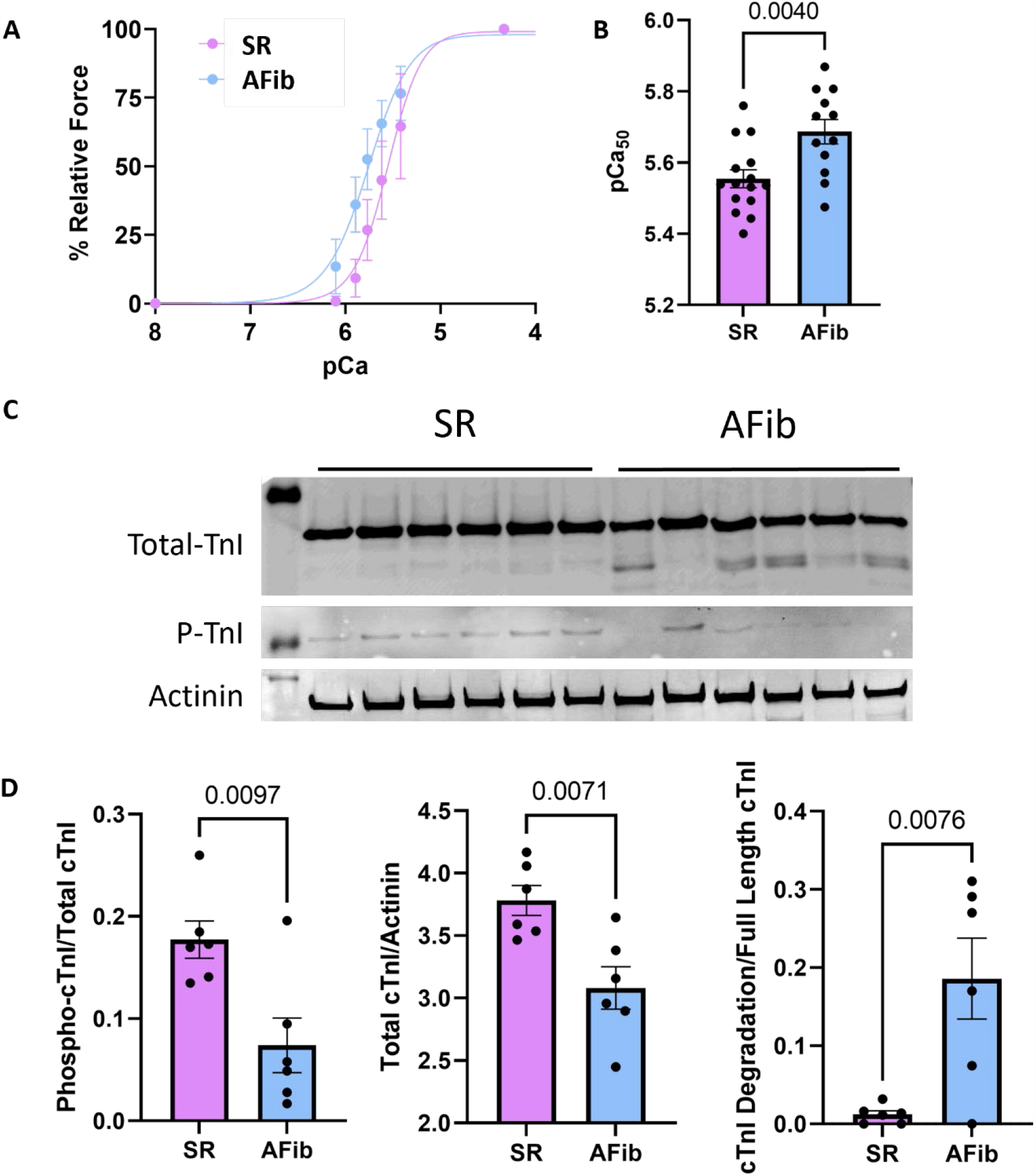
Atrial Fibrillation causes increased calcium sensitivity and TnI degradation. A) Skinned myocyte force-calcium relationship normalized to max force between SR and AFib hearts shows an increase in calcium sensitivity. B) Summary data for pCa50. D) Troponin I immunoblots. E) Analysis of TnI blots show a decrease in TnI phosphorylation and total TnI expression and an increase in TnI degradation.

### Atrial Fibrillation causes sensitization of calcium sensitivity of force development

After measuring maximum force, calcium sensitivity of force development was analyzed by normalizing the force of each cardiomyocyte to the maximum force of that cardiomyocyte. When normalized, calcium concentration can be analyzed against the percent relative force (Fig. 6A) which allows for determination of pCa50, the calcium concentration at half maximal force (Fig. 6B). The AFib atrial cardiomyocytes showed a significant increase in the pCa50 correlating to an increase in calcium sensitivity. We hypothesized that there would be a decrease in TnI phosphorylation in the AFib samples explains the increase in calcium sensitivity because, increased TnI phosphorylation at serine 23 and 24 can decrease calcium sensitivity (29). To test this hypothesis, immunoblots were run to analyze total TnI versus phosphorylated TnI (P-TnI) (Fig. 6C). Analysis of the immunoblot showed a decrease in the percent of phosphorylation between the AFib and SR groups and showed total TnI degradation, likely due to increased calpain targeted TnI degradation (Fig. 6D). Increased cTnI degradation and reduced cTnI phosphorylation of the full-length cTnI correlates with the increased calcium sensitivity of force development.

## (vi) DISCUSSION

Although AFib is characterized by electrophysiological dysfunction of the atria, the effect of AFib on atrial contractility is not well studied. We demonstrate that changes in sarcomere proteins and contractile deficits occur in AFib. We found that increased calpain activation in AFib correlates with widespread myofilament degradation including degradation of MHC, actinin, cMyBP-C, TnI, and potentially titin. The maximum force of contraction is decreased in AFib correlating to MHC degradation. Resting tension of the atrial cardiomyocytes is decreased in AFib correlating with titin degradation. Lastly, we found that calcium sensitivity is increased in AFib correlating to TnI cleavage and decreased TnI phosphorylation.

### Molecular alterations of myofilament proteins

The high amount of protein cleavage in this study makes interpreting the biophysical changes difficult. However, isolation of myofilament fractions enriched for intact sarcomere proteins suggests that some atrial myocytes were significantly degraded while others were functionally competent. The many biophysical changes we observed are consistent with our findings of specific protein functional effects. Even in intact sarcomeres, myofilament protein degradation was seen. It has been reported previously that there is a decrease in both the sarcomere content and the maximum force of contraction in AFib (30). The reduction in force generation was highly correlated with decreased sarcomere content, where patients with less sarcomere content also had a reduced force of contraction.

Increased calcium sensitivity may be explained by several molecular changes occurring in the atrial sarcomeres. The observed reduction in TnI phosphorylation from β-adrenergic activation of protein kinase A is well known to increase the calcium sensitivity of the myofilament (31). In AFib, it has been shown that there is an increase in PKA-dependent phosphorylation of calcium-handling proteins including ryanodine receptor, phospholamban, and the L-type calcium channel (32, 33). These results suggest that PKA activity is enhanced in AFib.

However, this increase in PKA-dependent phosphorylation has not been seen in myofilament proteins. Specifically, it has been shown that PKA-dependent phosphorylation of cMyBPC at serine 282 is decreased in AFib (33). With this data, it was suggested that activation of PKA in AFib is compartmentalized which produces hyper-phosphorylation of calcium handling proteins and hypo-phosphorylation of thick filament proteins (33). Our results suggest that this hypo-phosphorylation of myofilament proteins may extend to thin filament proteins as well as we have shown decreased TnI serine 23/24 phosphorylation. This theory of compartmentalized PKA activation in AFib may suggest that our current therapeutics that target β-adrenergic signaling and thus PKA activation, although effective in correcting calcium regulation in AFib, may be ineffective in correcting alterations within in the myofilament and thus ineffective for correcting contraction and proper blood ejection.

### Calpain proteolysis of sarcomere proteins

Calpain is a family of proteinases activated by high intracellular calcium (34). Once activated, calpain cleaves proteins at specific residues. This results in distinct cleavage patterns of its targets, rather than complete fragmentation of substrates (35). In cardiomyocytes, the calpain family of proteins is known to target a wide range of proteins including proteins involved in the cytoskeleton, apoptosis, signal transduction, calcium channels, and the myofilament (35, 36). Within the myofilament, calpain cleavage has been suggested to disconnect the proteins from the myofilament for them to be subsequently degraded by the proteasome, as it has been shown that intact myofibrils cannot be degraded by the proteasome (37).

Calpain activation and upregulation has been implicated in several cardiovascular diseases including myocardial infarction, ischemia and reperfusion, congestive heart failure, atrophy, DCM, and AFib (38-44). AFib results in cytosolic calcium overload resulting in increased calpain activity (45-48). Calpain activity in AFib has been correlated with both increased apoptosis and myofilament destruction (46, 48, 49). Calpain inhibition corrects the increase in apoptosis of the atrial myocytes in AFib (46). In this study, we show both increased calpain activity in AFib and the degradation of several calpain targets in the myofilament, expanding our understanding of calpain-mediated sarcomere destruction in AFib to include actinin and cMyBP-C as well as the contractile deficits that occur with these changes.

In addition to the full-length titin isoforms, there is an isoform called Cronos that is initiated from an intronic start site and includes the A-band region of titin with a small portion of the A/I junction. This Cronos titin, ∼2.2 MDa, is similar in size to the T2 product of titin, ∼2.3 MDa, which is thought to be a degradation product (50). Because we were unable to sufficiently separate the T2 and Cronos bands within the titin gels, we are unable to explicitly define the isoform increasing in T2/Cronos levels. As this is not a chronic model of AFib, and because we observed increased degradation of other myofilament proteins, it is parsimonious that the increase in the T2/Cronos band in the AFib samples is due to increased titin degradation and not increased Cronos expression. However, increasing either Cronos expression or titin proteolysis would account for the observed reduction in resting tension in the AFib cardiomyocytes. To our knowledge, the presence of Cronos in chronic AFib has not been explored.

### The role of contractility in thrombus development and stroke risk in AFib

As the risk for thrombus development is high in atrial fibrillation, understanding the aspects of this riskis essential for both the management of AFib and its prevention. Thrombus development leads to increased risk of stroke development in AFib. Myocardial thrombus development is a result of myocardial injury, reduced wall motion, and reduced blood flow that are key aspects of thrombus development but remain incompletely understood in AFib. While these aspects may, in part, be described by alterations in the electrical signal propagation throughout the atria leading to improper excitation, the role of impaired contractility in maintaining blood flow through the atria remains under investigated in AFib and its progression. Recently, computational modeling of healthy atrial and AFib atrial contractility based on the wall motion, wall velocity, and morphology of the left atrial appendage indicates that these parameters result in reduced shear strain rate, and thus impaired contractility in AFib and is associated with increased risk for thrombus development. This highlights that there are underlying alterations in contractility within the AFib that is yet to be understood.

### Atrial stunning and contractile recovery following rhythm restoration

AFib is currently treated using atrial cardioversion (51). The goal of these treatments is to restore the normal rhythm of the atria. It has long been noted that after cardioversion and successful restoration of atrial rhythm, the atria experience a period of mechanical dysfunction known as atrial stunning (10, 12, 13). Atrial stunning may be resolved within minutes of cardioversion or take up to 4 – 6 weeks, with recovery duration generally correlating with left atrial size, co-morbid structural heart diseases, and the duration of the preceding AFib (15, (12, 13). Within this period of stunning, patients are at an increased risk for thromboembolic events including stroke (10, 52). The management of atrial cardioversion and atrial stunning has been progressing in recent years with the use of transesophageal echocardiogram directed cardioversion (52). Although management during atrial stunning has been improving, there is still little understanding of the pathology of atrial stunning and thus prevention of atrial stunning remains incomplete.

Atrial stunning was thought to be a product of the electrical current delivered to the atrial myocardium (10). It was later found that atrial stunning also occurs in pharmacological cardioversion and does not occur after electrical current delivery to healthy myocardium (53, 54). Therefore, atrial stunning is at least partially a consequence of AFib rather than a product of cardioversion therapy (55). Although this phenomenon has been widely known for decades, there is little understanding of the molecular mechanisms behind atrial stunning (15, 45, 46). There are currently three main suggested mechanisms behind atrial stunning including tachycardia-induced atrial cardiomyopathy, cytosolic calcium overload, and atrial dedifferentiation (10). Cytosolic calcium overload has been thought to contribute to atrial stunning via the L-type calcium channel and it was found that BayY5959, an L-type calcium channel agonist, partially restored atrial contractility after a 3-day duration of AFib (56). This suggests that although calcium overload may play a role in atrial stunning, correction of this calcium overload is not completely effective in restoring atrial contractility and thus preventing atrial stunning.

Our data suggest a molecular mechanism that may contribute to atrial stunning. We have shown that wide-spread myofilament proteolysis occurs in AFib. Further investigation of this would require defining whether the period of atrial stunning depends on removal of degraded proteins and synthesis of new full-length proteins to restore the integrity of the myofilaments to a sufficient degree for functional recovery. Determining whether increased duration of sustained AFib corresponds with increased myofilament proteolysis also remains to be studied. Defining this molecular mechanism in atrial stunning will provide a target for treatment of stunning post-cardioversion treatment of AFib.

## (xiv) SUPPLEMENTARY DATA

Figures S1 – S4 and Table S1 and S2 are found in the supplementary data.

## (vii) FUNDING

This work has been supported by NIH grants NHLBI R00141698, R56165137 (DYB), 1R35HL161249-01, and American Heart Association grant 23POST1023125 (AAA).

## (x) CONFLICT OF INTEREST

The authors have no conflicts of interest regarding this work.

## (viii) AUTHOR CONTRIBUTIONS

H.E.C., R.A., and D.Y.B. conceived and designed research; H.E.C., H.V.B., A.P., A.A.A., T.D., C.D., K.A., and S.E. performed experiments; H.E.C., H.V.B., A.P., A.A.A., T.D., C.D., S.E., and D.Y.B. analyzed data; H.E.C., H.V.B., J.A.K., R.A., and D.Y.B. interpreted results of experiments; H.E.C. and D.Y.B. prepared figures; H.E.C., S.E., and D.Y.B. drafted the manuscript; H.E.C., J.A.K., R.A., and D.Y.B. edited and revised the manuscript; all authors approved the final version of the manuscript.

## ABBREVIATIONS

(AFib): Atrial fibrillation
(SR): sinus rhythm
(RAP): rapid atrial pacing
(GO): gene ontology
(MHC): myosin heavy chain
(cMyBP-C): cardiac myosin binding protein-C
(cTnI): cardiac Troponin I
(SRX): super relaxed state of myosin

