## Supplement for "Proteolytic degradation of atrial sarcomere proteins underlies contractile defects in atrial fibrillation"

### SUPPLEMENTARY TABLES/FIGURES

| **All Significantly Downregulated Genes** | | | | | |
|---|---|---|---|---|---|
| Gene | Description | AFib Average | SR Average | Abundance Ratio: (AFib) / (SR) | Abundance Ratio P-Value: (AFib) / (SR) |
| ADAMTSL1 | ADAMTS like 1 OS | 0 | 1178700.869 | 0.01 | 1E-17 |
| CACNB2 | Calcium voltage-gated channel auxiliary subunit beta 2 OS | 0 | 317606.5978 | 0.01 | 1E-17 |
| CILP2 | Cartilage intermediate layer protein 2 OS | 0 | 921198.6913 | 0.01 | 1E-17 |
| CSDC2 | Cold shock domain containing C2 OS | 0 | 201667.1651 | 0.01 | 1E-17 |
| FERMT2 | FERM domain containing kindlin 2 OS | 0 | 246338.9309 | 0.01 | 1E-17 |
| MLIP | Muscular LMNA interacting protein OS | 0 | 552509.9459 | 0.01 | 1E-17 |
| MYH7B | Myosin heavy chain 7B OS | 0 | 236888.5329 | 0.01 | 1E-17 |
| PKP1 | Plakophilin 1 OS | 0 | 610765.1134 | 0.01 | 1E-17 |
| SMTNL2 | Uncharacterized protein OS | 0 | 614785.9443 | 0.01 | 1E-17 |
| UBE2D1 | Ubiquitin conjugating enzyme E2 D1 OS | 0 | 254473.2277 | 0.01 | 1E-17 |
| YBX3 | Y-box binding protein 3 OS | 0 | 335002.0341 | 0.01 | 1E-17 |
| KRT31 | Uncharacterized protein OS | 131846.6531 | 50673331.42 | 0.013 | 1E-17 |
| KRT35 | Keratin 35 OS | 146008.5625 | 2801486.673 | 0.07 | 1.42109E-14 |
| ARF1 | ADP ribosylation factor 1 OS | 238097.4852 | 1783359.175 | 0.128 | 4.19612E-09 |
| COL4A1 | Collagen type IV alpha 1 chain OS | 3724854.654 | 31062074.62 | 0.2 | 2.44268E-05 |
| PPCS | Phosphopantothenoylcysteine synthetase OS | 90175.20313 | 447114.834 | 0.202 | 0.000590799 |
| PBXIP1 | PBX homeobox interacting protein 1 OS | 140104.4592 | 864871.981 | 0.213 | 8.4971E-06 |
| TNNC1 | Troponin C1, slow skeletal and cardiac type OS | 1574150.362 | 5801752.751 | 0.213 | 3.66924E-05 |
| COL4A2 | Collagen type IV alpha 2 chain OS | 3052251.247 | 17965164.94 | 0.222 | 7.83927E-05 |
| RNF7 | Ring finger protein 7 OS | 225688.3842 | 989831.389 | 0.228 | 7.32027E-06 |
| TUBB1 | Tubulin beta 1 class VI OS | 122417.5858 | 539459.454 | 0.231 | 0.000157772 |
| FAM180B | Family with sequence similarity 180 member B OS | 25058.76591 | 68760.44503 | 0.232 | 0.002489906 |
| S100A1 | S100 calcium binding protein A1 OS | 637986.4049 | 3657070.436 | 0.251 | 7.33405E-05 |
| MMP3 | Stromelysin-1 OS | 90031.08149 | 357072.4202 | 0.252 | 0.002402432 |
| FHL3 | Four and a half LIM domains 3 OS | 166105.4363 | 620531.3683 | 0.268 | 0.000333388 |
| FBXW11 | F-box and WD repeat domain containing 11 OS | 71267.25724 | 262806.8809 | 0.271 | 0.00297561 |
| DSG1 | Desmoglein-1 OS | 210106.3826 | 289346.8371 | 0.278 | 0.003442865 |
| DPY19L1 | Dpy-19 like C-mannosyltransferase 1 OS | 137977.4032 | 475233.8103 | 0.29 | 0.00226428 |
| CDIN1 | CDAN1 interacting nuclease 1 OS | 783813.2662 | 2266123.616 | 0.294 | 0.000192977 |
| OLFML3 | Olfactomedin like 3 OS | 645125.2613 | 2108367.957 | 0.306 | 0.000330259 |
| PTMS | Parathymosin OS | 166878.0085 | 713004.9252 | 0.311 | 0.000687198 |
| ADSS1 | Adenylosuccinate synthase 1 OS | 286456.3466 | 2426195.694 | 0.315 | 0.000414856 |
| SNAP47 | Uncharacterized protein OS | 14830.16482 | 46083.80476 | 0.322 | 0.019658392 |
| TOMM20 | Translocase of outer mitochondrial membrane 20 OS | 28258.72062 | 262092.0658 | 0.332 | 0.009673045 |
| LNPK | Lunapark, ER junction formation factor OS | 259881.8678 | 313669.2678 | 0.339 | 0.012787151 |
| COL6A6 | Collagen type VI alpha 6 chain OS | 1457704.112 | 5943689.811 | 0.34 | 0.003451308 |
| CES2 | Carboxylesterase 2 OS | 843657.4236 | 2286095.784 | 0.342 | 0.000882774 |
| CYP27A1 | Cytochrome P450 family 27 subfamily A member 1 OS | 295551.0522 | 752313.9931 | 0.347 | 0.001248718 |
| MOGAT1 | Monoacylglycerol O-acyltransferase 1 OS | 703636.2968 | 2191583.104 | 0.348 | 0.001474741 |
| CUL4B | Cullin 4B OS | 235274.4734 | 675104.0365 | 0.349 | 0.001617271 |
| TPM2 | Tropomyosin 2 OS | 2698065.111 | 7710866.91 | 0.35 | 0.005400103 |
| ASS1 | Argininosuccinate synthase 1 OS | 35182.38749 | 100240.9131 | 0.351 | 0.018632097 |
| NPC1 | NPC intracellular cholesterol transporter 1 OS | 195650.4639 | 436487.7168 | 0.361 | 0.009270258 |
| ANK1 | Ankyrin 1 OS | 689712.7279 | 2077757.098 | 0.365 | 0.002094607 |
| GPD1 | Glycerol-3-phosphate dehydrogenase 1 OS | 4434588.915 | 11824622.26 | 0.371 | 0.008251444 |
| POMGNT2 | Protein O-linked-mannose beta-1,4-N-acetylglucosaminyltransferase 2 OS | 184654.1849 | 437316.5868 | 0.373 | 0.01129443 |
| PRDM12 | PR/SET domain 12 OS | 16211904.8 | 40036196.27 | 0.381 | 0.010092639 |
| YIF1A | Yip1 interacting factor homolog A, membrane trafficking protein OS | 238325.5498 | 844775.8339 | 0.382 | 0.003739983 |
| MRPL12 | Mitochondrial ribosomal protein L12 OS | 44766.63277 | 106208.2504 | 0.384 | 0.029547402 |
| MYLK3 | Myosin light chain kinase 3 OS | 1929343.105 | 3586629.766 | 0.386 | 0.008170665 |
| ADAMTSL5 | Uncharacterized protein OS | 278165.5692 | 604769.5663 | 0.393 | 0.005755449 |
| LOC482182 | Uncharacterized protein OS | 961670.5289 | 3557058.218 | 0.394 | 0.007313206 |
| PACS1 | Phosphofurin acidic cluster sorting protein 1 OS | 237979.2509 | 600160.0772 | 0.397 | 0.010237724 |
| HYAL1 | Uncharacterized protein OS | 221132.4453 | 385407.0565 | 0.399 | 0.023169524 |
| MVB12B | Multivesicular body subunit 12B OS | 213683.5169 | 470874.2562 | 0.4 | 0.013038648 |
| NEFH | Uncharacterized protein OS | 185350.3891 | 462925.2828 | 0.4 | 0.017597159 |
| NT5DC3 | 5'-nucleotidase domain containing 3 OS | 216220.1098 | 689383.8934 | 0.4 | 0.006462525 |
| CMSS1 | Collagen type VIII alpha 1 chain OS | 128530.6525 | 1139834.043 | 0.401 | 0.004496857 |
| NDUFAB1 | Uncharacterized protein OS | 1206835.557 | 2966958.567 | 0.402 | 0.007855761 |
| SIAE | Sialic acid acetylesterase OS | 265368.9506 | 571551.9477 | 0.404 | 0.011780717 |

| | | | | | |
|---|---|---|---|---|---|
| HSD17B13 | Hydroxysteroid 17-beta dehydrogenase 11 OS | 3475185.909 | 10025595.39 | 0.405 | 0.015397132 |
| OLFML1 | Synaptotagmin 9 OS | 201395.1563 | 380679.3379 | 0.406 | 0.034934659 |
| COQ4 | Coenzyme Q4 OS | 3645552.032 | 7532294.954 | 0.416 | 0.018673199 |
| OCIAD1 | OCIA domain containing 1 OS | 1006609.636 | 2026466.313 | 0.416 | 0.007325603 |
| ETFRF1 | Electron transfer flavoprotein regulatory factor 1 OS | 758382.5743 | 1374732.503 | 0.417 | 0.008285589 |
| LTBP1 | Latent transforming growth factor beta binding protein 1 OS | 232356.1313 | 1186588.32 | 0.421 | 0.007419851 |
| EMC10 | ER membrane protein complex subunit 10 OS | 242561.6514 | 562304.5167 | 0.431 | 0.017733713 |
| MPC1 | Mitochondrial pyruvate carrier 1 OS | 290567.9849 | 579169.2227 | 0.432 | 0.016534118 |
| CA14 | Carbonic anhydrase 14 OS | 141459.3245 | 752784.7842 | 0.436 | 0.013621583 |
| USP22 | Uncharacterized protein OS | 504354.9488 | 1205792.346 | 0.438 | 0.013177809 |
| 4 | Uncharacterized protein OS | 709687.8868 | 3714821.537 | 0.444 | 0.019307359 |
| SPON1 | Spondin 1 OS | 73323.36389 | 633000.6323 | 0.444 | 0.023444493 |
| COX16 | Cytochrome c oxidase assembly factor COX16 OS | 2055547.884 | 4695724.54 | 0.445 | 0.022710071 |
| PIN4 | Peptidylprolyl cis/trans isomerase, NIMA-interacting 4 OS | 430656.796 | 946753.5233 | 0.447 | 0.012346462 |
| PCOLCE | Procollagen C-endopeptidase enhancer OS | 526725.1146 | 1248548.326 | 0.448 | 0.015484072 |
| MFAP4 | Microfibril associated protein 4 OS | 53200338.17 | 96107180.42 | 0.45 | 0.031368302 |
| DHRS7C | Dehydrogenase/reductase 7C OS | 441670.9285 | 908426.907 | 0.451 | 0.012598328 |
| ARPIN | Actin related protein 2/3 complex inhibitor OS | 817699.5016 | 1795408.66 | 0.455 | 0.015933839 |
| PRMT7 | Protein arginine methyltransferase 7 OS | 271309.0561 | 367282.1738 | 0.455 | 0.038505901 |
| CNIH4 | Cornichon family AMPA receptor auxiliary protein 4 OS | 1016584.076 | 2172963.526 | 0.456 | 0.012952825 |
| RAB9B | RAB9B, member RAS oncogene family OS | 337570.818 | 722494.3882 | 0.457 | 0.015571423 |
| ALPK3 | Alpha kinase 3 OS | 339839.7685 | 495879.6344 | 0.459 | 0.027516082 |
| MLF1 | Myeloid leukemia factor 1 OS | 198197.6579 | 414938.1143 | 0.459 | 0.047756815 |
| EMILIN1 | Elastin microfibril interfacer 1 OS | 814814.6961 | 1667565.353 | 0.46 | 0.019035321 |
| LOC474938 | Uncharacterized protein OS | 2895187.76 | 4576025.32 | 0.461 | 0.033245478 |
| LRBA | LPS responsive beige-like anchor protein OS | 425738.3098 | 922595.0368 | 0.461 | 0.016490983 |
| ITM2B | Integral membrane protein 2B OS | 1152989.898 | 248698.7515 | 0.462 | 0.016155122 |
| DTD1 | Uncharacterized protein OS | 420007.5538 | 1087774.541 | 0.465 | 0.019640844 |
| TNNI1 | Troponin I1, slow skeletal type OS | 403968.5579 | 931915.604 | 0.471 | 0.021250841 |
| CRYM | Uncharacterized protein OS | 18716524.2 | 43116314.18 | 0.473 | 0.042861626 |
| ISG15 | ISG15 ubiquitin like modifier OS | 348582.5473 | 812483.5484 | 0.473 | 0.017559129 |
| CNTFR | Ciliary neurotrophic factor receptor subunit alpha OS | 341555.1687 | 632004.672 | 0.476 | 0.020987765 |
| HSD17B13 | Hydroxysteroid 17-beta dehydrogenase 11 OS | 905101.9194 | 1726691.19 | 0.477 | 0.022712296 |
| UQCRH | Uncharacterized protein OS | 6547225.769 | 13127753.21 | 0.479 | 0.04661218 |
| ART3 | ADP-ribosyltransferase 3 (inactive) OS | 884277.8578 | 1843420.453 | 0.48 | 0.021490816 |
| NDRG2 | NDRG family member 2 OS | 5758877.626 | 11340916.98 | 0.48 | 0.046809476 |
| ADAMTSL4 | ADAMTS like 4 OS | 438975.3587 | 837385.0017 | 0.482 | 0.020241693 |
| GNB3 | Guanine nucleotide-binding protein G(I)/G(S)/G(T) subunit beta-3 OS | 214260.0666 | 451936.3841 | 0.482 | 0.046368794 |
| LDB3 | LIM domain binding 3 OS | 2762114.797 | 6729974.11 | 0.482 | 0.044081975 |
| TRPT1 | tRNA phosphotransferase 1 OS | 264832.2693 | 549178.2464 | 0.482 | 0.038579313 |
| UBE2K | Ubiquitin conjugating enzyme E2 K OS | 772519.2419 | 1845116.473 | 0.485 | 0.02595119 |
| VAC14 | VAC14 component of PIKFYVE complex OS | 701364.3034 | 1030910.435 | 0.485 | 0.027556913 |
| CDIPT | Uncharacterized protein OS | 371776.228 | 913270.0591 | 0.486 | 0.019656008 |
| KRT18 | Keratin 18 OS | 805473.7566 | 1653848.791 | 0.487 | 0.028750545 |
| ASL | Uncharacterized protein OS | 1783999.876 | 3499918.506 | 0.488 | 0.03589736 |
| TOMM22 | Uncharacterized protein OS | 1121182.184 | 2137038.998 | 0.489 | 0.023296833 |
| MRPL23 | Uncharacterized protein OS | 1086995.947 | 2358694.431 | 0.492 | 0.022483552 |
| PELO | Integrin subunit alpha 1 OS | 1567818.76 | 4376056.105 | 0.494 | 0.048505583 |
| STAU2 | Staufen double-stranded RNA binding protein 2 OS | 1032935.867 | 1950936.125 | 0.494 | 0.028939474 |

***Supplemental Table 1:*** *Significantly downregulated genes in Atrial Fibrillation*

| **All Significantly Upregulated Genes** | | | | | |
|---|---|---|---|---|---|
| Gene | Description | AFib Average | SR Average | Abundance Ratio: (AFib) / (SR) | Abundance Ratio P-Value: (AFib) / (SR) |
| PIKFYVE | Phosphoinositide kinase, FYVE-type zinc finger containing OS | 2085238.021 | 991826.7397 | 2.102 | 0.045010401 |
| CFD | Uncharacterized protein OS | 744365.8843 | 461258.7077 | 2.183 | 0.043102025 |
| APOE | Apolipoprotein E OS | 3751258.622 | 946688.0903 | 2.194 | 0.048143734 |
| MAP7D1 | Uncharacterized protein OS | 763273.9047 | 304290.4759 | 2.196 | 0.044977754 |
| ENSA | Endosulfine alpha OS | 1756669.361 | 746109.1861 | 2.198 | 0.040882582 |
| ITIH2 | Inter-alpha-trypsin inhibitor heavy chain 2 OS | 2492857.587 | 1175701.01 | 2.202 | 0.038274386 |

| | | | | | |
|---|---|---|---|---|---|
| RBBP4 | RB binding protein 4, chromatin remodeling factor OS | 1568049.397 | 785350.9001 | 2.205 | 0.037402498 |
| BIN2 | Bridging integrator 2 OS | 1016510.324 | 206970.121 | 2.206 | 0.042112771 |
| PLA2G7 | Platelet-activating factor acetylhydrolase OS | 1222028.272 | 356146.3265 | 2.214 | 0.046435697 |
| RPS27L | Ribosomal protein S27 like OS | 1726212.644 | 580547.9062 | 2.225 | 0.032434942 |
| CKAP4 | Cytoskeleton associated protein 4 OS | 4936825.67 | 1644804.885 | 2.234 | 0.047713282 |
| ELMO1 | Engulfment and cell motility 1 OS | 957793.8183 | 370705.9835 | 2.245 | 0.045176653 |
| ATG3 | Autophagy related 3 OS | 668689.5745 | 613465.7452 | 2.25 | 0.038783621 |
| ACTR3B | Uncharacterized protein OS | 2558693.829 | 461356.8395 | 2.26 | 0.030291453 |
| PRKRA | Uncharacterized protein OS | 1172552.668 | 586124.9476 | 2.261 | 0.043511415 |
| MICU2 | Mitochondrial calcium uptake 2 OS | 966852.783 | 473745.3404 | 2.262 | 0.0494773 |
| RPS17 | 40S ribosomal protein S17 OS | 3540153.47 | 2265681.82 | 2.266 | 0.043920443 |
| F2 | Coagulation factor II, thrombin OS | 5401851.624 | 1261376.091 | 2.268 | 0.041661541 |
| CFL1 | Cofilin 1 OS | 20088781.53 | 15156189.69 | 2.269 | 0.048546974 |
| LOC477072 | Uncharacterized protein OS | 80561429.9 | 29925861.32 | 2.289 | 0.045997212 |
| H3C4 | H3 clustered histone 4 OS | 2213605.552 | 1003121.372 | 2.294 | 0.027822829 |
| ABRACL | ABRA C-terminal like OS | 2536098.27 | 1664634.613 | 2.304 | 0.034792653 |
| PGRMC1 | Progesterone receptor membrane component 1 OS | 3121527.834 | 1354304.341 | 2.305 | 0.042131942 |
| S100A9 | Uncharacterized protein OS | 2159470.467 | 571228.2384 | 2.309 | 0.024291177 |
| TRIM55 | Tripartite motif containing 55 OS | 8852651.114 | 3261668.098 | 2.337 | 0.040589971 |
| SPO11 | SPO11 initiator of meiotic double stranded breaks OS | 691677.0716 | 295892.897 | 2.338 | 0.03685163 |
| KNG1 | Kininogen 1 OS | 8544704.87 | 2799042.038 | 2.353 | 0.038873099 |
| SPARCL1 | SPARC like 1 OS | 1867654.152 | 351166.2613 | 2.353 | 0.019007921 |
| CFH | Uncharacterized protein OS | 10570901.68 | 3007293.745 | 2.358 | 0.03838787 |
| BLVRB | Biliverdin reductase B OS | 766756.0619 | 287209.0473 | 2.359 | 0.029226466 |
| C5 | Complement C5 OS | 1521794.877 | 207012.8637 | 2.36 | 0.035875333 |
| CA1 | Carbonic anhydrase 1 OS | 8664436.042 | 2268607.497 | 2.371 | 0.037048684 |
| TIMM21 | Translocase of inner mitochondrial membrane 21 OS | 2158578.562 | 758765.6861 | 2.378 | 0.021413792 |
| PTPRC | Uncharacterized protein OS | 1588327.586 | 447443.2785 | 2.381 | 0.022941081 |
| KRT9 | Keratin, type I cytoskeletal 9 OS | 794999.5597 | 378111.8254 | 2.4 | 0.021227549 |
| HSPH1 | Heat shock protein family H (Hsp110) member 1 OS | 1230523.187 | 751525.3472 | 2.404 | 0.024416581 |
| MAP1A | Microtubule associated protein 1A OS | 942640.001 | 458439.6985 | 2.408 | 0.026463969 |
| LOC119881797 | Uncharacterized protein OS | 808131.3125 | 334921.3868 | 2.413 | 0.024153639 |
| RPS20 | Uncharacterized protein OS | 2585627.202 | 673833.1931 | 2.417 | 0.018049718 |
| GPX4 | Glutathione peroxidase 4 OS | 1827677.219 | 765433.3603 | 2.42 | 0.019841627 |
| MMP9 | Matrix metalloproteinase-9 OS | 6350483.834 | 230093.2682 | 2.421 | 0.029065426 |
| PRMT1 | Uncharacterized protein OS | 1157608.614 | 575308.7397 | 2.426 | 0.028226049 |
| LYRM7 | Uncharacterized protein OS | 2641777.216 | 1434168.884 | 2.443 | 0.023134518 |
| CTSZ | Cathepsin Z OS | 1390419.949 | 451656.5572 | 2.452 | 0.03063734 |
| RPL30 | Uncharacterized protein OS | 1503984.592 | 362171.6623 | 2.453 | 0.029837189 |
| CCN2 | Cellular communication network factor 2 OS | 1393478.68 | 454373.3201 | 2.457 | 0.030207768 |
| MAPKAPK2 | MAPK activated protein kinase 2 OS | 1428971.359 | 468640.1956 | 2.461 | 0.026946379 |
| GCHFR | GTP cyclohydrolase I feedback regulator OS | 3872260.912 | 788583.0837 | 2.462 | 0.021996179 |
| MAT2B | Uncharacterized protein OS | 634483.5649 | 257459.3767 | 2.464 | 0.030275367 |
| JAG2 | Jagged canonical Notch ligand 2 OS | 1302493.014 | 249494.4604 | 2.47 | 0.023603808 |
| SERPING1 | Serpin family G member 1 OS | 2612984.57 | 774789.099 | 2.47 | 0.015955479 |
| UBE2M | Ubiquitin conjugating enzyme E2 M OS | 809050.054 | 124269.3471 | 2.487 | 0.028235066 |
| AGT | Angiotensinogen OS | 2071906.255 | 738795.9709 | 2.497 | 0.014598028 |
| RPS19 | Uncharacterized protein OS | 4330422.135 | 592708.0978 | 2.506 | 0.018092591 |
| THBS1 | Thrombospondin 1 OS | 1234165.226 | 151974.0306 | 2.512 | 0.020191633 |
| PSMA3 | Proteasome 20S subunit alpha 3 OS | 1157741.615 | 1632181.158 | 2.524 | 0.013570383 |
| ACOX1 | Acyl-CoA oxidase 1 OS | 1127298.389 | 257812.8473 | 2.526 | 0.019727717 |
| COTL1 | Coactosin like F-actin binding protein 1 OS | 891661.3941 | 250366.5455 | 2.531 | 0.017868529 |
| LGALS3 | Galectin-3 OS | 2761200.497 | 1153039.109 | 2.558 | 0.015774208 |
| PADI2 | Peptidyl arginine deiminase 2 OS | 1092656.306 | 142656.952 | 2.561 | 0.017136956 |
| TOMM40 | Translocase of outer mitochondrial membrane 40 OS | 1634698.089 | 598687.807 | 2.561 | 0.008640592 |
| MGAM | Maltase-glucoamylase OS | 970042.7812 | 119476.1837 | 2.566 | 0.014856389 |
| PREB | Prolactin regulatory element binding OS | 691606.5941 | 308615.6365 | 2.568 | 0.017886944 |
| LOC100685620 | Uncharacterized protein OS | 8136048.862 | 3301190.346 | 2.578 | 0.021571202 |
| SERPINC1 | Serpin family C member 1 OS | 11194596.3 | 4659624.928 | 2.582 | 0.021322445 |
| CRK | CRK proto-oncogene, adaptor protein OS | 1102319.814 | 445828.4721 | 2.587 | 0.016821203 |
| CD44 | CD44 antigen OS | 5231757.266 | 1695522.511 | 2.59 | 0.017655079 |
| MRPS24 | Uncharacterized protein OS | 555087.7856 | 260947.3412 | 2.593 | 0.030882345 |
| HRG | Histidine rich glycoprotein OS | 1718741.58 | 469301.158 | 2.595 | 0.010651238 |
| KRT80 | Keratin 80 OS | 1180145.247 | 453796.5124 | 2.601 | 0.017402953 |
| GAK | Cyclin G associated kinase OS | 482848.4185 | 238114.0769 | 2.612 | 0.034979982 |
| CLTA | Clathrin light chain A OS | 609120.2187 | 232907.2769 | 2.615 | 0.029445432 |
| SNRNP70 | Small nuclear ribonucleoprotein U1 subunit 70 OS | 509067.9061 | 218196.52 | 2.616 | 0.03414173 |
| RPL27 | 60S ribosomal protein L27 OS | 5314197.783 | 3574603.994 | 2.624 | 0.02081525 |
| RAD23A | RAD23 homolog A, nucleotide excision repair protein OS | 3180003.25 | 1210979.338 | 2.626 | 0.017897075 |
| SLC25A5 | Uncharacterized protein OS | 4946218.681 | 1453395.719 | 2.633 | 0.016722363 |
| F13A1 | Coagulation factor XIII A chain OS | 2563609.913 | 856052.6309 | 2.637 | 0.008907276 |
| CP | Ceruloplasmin OS | 5220489.193 | 1171683.491 | 2.676 | 0.014990393 |

| | | | | | |
|---|---|---|---|---|---|
| ITGAD | Integrin subunit alpha D OS | 1388604.02 | 185522.377 | 2.682 | 0.011415767 |
| SHBG | Sex hormone binding globulin OS | 933131.0421 | 261236.726 | 2.689 | 0.009157094 |
| ARPP19 | Uncharacterized protein OS | 1381628.277 | 695498.0063 | 2.693 | 0.007390007 |
| LETMD1 | LETM1 domain containing 1 OS | 399687.3546 | 206525.5524 | 2.694 | 0.039226937 |
| LOC119880472 | FKBP prolyl isomerase 1B OS | 693750.6807 | 296148.0364 | 2.697 | 0.01286271 |
| CAP1 | Uncharacterized protein OS | 3263890.715 | 955244.9029 | 2.717 | 0.010248361 |
| RTN3 | Reticulon 3 OS | 178090.2767 | 78716.76355 | 2.718 | 0.031377197 |
| RPL24 | Ribosomal protein L24 OS | 7381797.91 | 2701964.125 | 2.732 | 0.014990281 |
| 607076 | Monocyte differentiation antigen CD14 OS | 330748.9033 | 73709.06826 | 2.741 | 0.047709561 |
| APOBEC2 | Apolipoprotein B mRNA editing enzyme catalytic subunit 2 OS | 22078827.15 | 9720138.909 | 2.777 | 0.012865767 |
| MYO1F | Myosin IF OS | 535349.189 | 27273.88822 | 2.788 | 0.041676635 |
| F13B | Coagulation factor XIII B chain OS | 383996.6042 | 6130.386399 | 2.798 | 0.040452767 |
| APOC1 | Apolipoprotein C-I OS | 6298382.471 | 790092.5037 | 2.806 | 0.0102804 |
| LOC119881719 | Ubiquitin conjugating enzyme E2 V1 OS | 610043.7282 | 187870.9697 | 2.83 | 0.016735491 |
| HABP2 | Hyaluronan binding protein 2 OS | 1625062.348 | 448719.7225 | 2.833 | 0.004856923 |
| PLEKHA5 | Pleckstrin homology domain containing A5 OS | 228049.625 | 124035.1104 | 2.84 | 0.037653007 |
| AFM | Afamin OS | 2309991.06 | 499996.5085 | 2.843 | 0.004893804 |
| ACTBL2 | Actin beta like 2 OS | 51250042.22 | 12901711.46 | 2.847 | 0.01072531 |
| LMNA | Lamin A/C OS | 13626393.13 | 2165419.466 | 2.851 | 0.010637967 |
| PCBD1 | Pterin-4 alpha-carbinolamine dehydratase 1 OS | 2471085.193 | 475964.9648 | 2.854 | 0.004834953 |
| RIPK1 | Receptor interacting serine/threonine kinase 1 OS | 163792.5266 | 75929.76996 | 2.877 | 0.025221422 |
| CAPG | Capping actin protein, gelsolin like OS | 2948042.461 | 531582.5894 | 2.923 | 0.003977455 |
| NUDT5 | Nudix hydrolase 5 OS | 1439604.307 | 488548.036 | 2.947 | 0.006615006 |
| VASP | Vasodilator-stimulated phosphoprotein OS | 2185932.674 | 492992.9448 | 2.985 | 0.002678798 |
| NEFM | Uncharacterized protein OS | 81472.93014 | 49280.57256 | 2.999 | 0.033993351 |
| SRI | Sorcin OS | 7783681.829 | 3121688.282 | 3.001 | 0.007232818 |
| TRADD | TNFRSF1A associated via death domain OS | 335422.8709 | 85935.9981 | 3.005 | 0.027909244 |
| HRC | Histidine rich calcium binding protein OS | 9889351.022 | 2940228.34 | 3.039 | 0.006577729 |
| PGLYRP1 | Peptidoglycan recognition protein 1 OS | 2352445.992 | 1043986.874 | 3.054 | 0.002419845 |
| LOC476006 | Uncharacterized protein OS | 7946046.191 | 2130321.045 | 3.057 | 0.00653296 |
| GC | GC vitamin D binding protein OS | 13105057.56 | 4288867.217 | 3.059 | 0.006250931 |
| CENPV | Uncharacterized protein OS | 543205.9944 | 177346.4438 | 3.063 | 0.012315824 |
| LOC477562 | Uncharacterized protein OS | 415711.9182 | 240604.5091 | 3.076 | 0.013469758 |
| COX6A2 | Cytochrome c oxidase subunit 6A2, mitochondrial OS | 359338.3114 | 14610.22561 | 3.098 | 0.023977122 |
| NPPA | Natriuretic peptides A OS | 4354743.239 | 1116596.614 | 3.109 | 0.004372737 |
| KRT15 | Keratin 15 OS | 9544266.054 | 1189789.863 | 3.114 | 0.005435344 |
| FN1 | Fibronectin OS | 24410629.58 | 5207834.327 | 3.128 | 0.005250651 |
| PON1 | Paraoxonase 1 OS | 961893.7644 | 169439.6447 | 3.135 | 0.002706242 |
| PFKP | Phosphofructokinase, platelet OS | 883775.4701 | 219082.7636 | 3.2 | 0.002101948 |
| BTNL9 | Butyrophilin like 9 OS | 367710.8065 | 37737.1646 | 3.202 | 0.020330889 |
| ARPC3 | Uncharacterized protein OS | 2801580.731 | 854295.7583 | 3.203 | 0.001978202 |
| ANKRD1 | Ankyrin repeat domain 1 OS | 5295841.11 | 1260915.978 | 3.233 | 0.003381475 |
| PABPC1L | Poly(A) binding protein cytoplasmic 1 like OS | 2644178.38 | 757490.1207 | 3.241 | 0.001353958 |
| TGM2 | Transglutaminase 2 OS | 47178595.13 | 11707088.02 | 3.252 | 0.003853715 |
| THEMIS2 | Thymocyte selection associated family member 2 OS | 183529.9297 | 31344.429 | 3.273 | 0.012647031 |
| EDF1 | Uncharacterized protein OS | 791442.5368 | 144520.7474 | 3.275 | 0.0032871 |
| CNN2 | Uncharacterized protein OS | 1865957.381 | 559569.4828 | 3.298 | 0.001049985 |
| TMEM109 | Transmembrane protein 109 OS | 4727660.151 | 1430069.045 | 3.306 | 0.002592852 |
| EFL1 | Elongation factor like GTPase 1 OS | 138225.8494 | 45635.19335 | 3.317 | 0.011488998 |
| HMGB2 | High mobility group box 2 OS | 3370831.803 | 1859995.778 | 3.332 | 0.002676673 |
| RABGAP1 | RAB GTPase activating protein 1 OS | 59908.29397 | 19958.65501 | 3.343 | 0.031161085 |
| CEACAM1 | CEA cell adhesion molecule 1 OS | 1142452.844 | 130288.8856 | 3.355 | 0.00129866 |
| H2BC15 | Histone H2B OS | 522488.1523 | 179368.4671 | 3.37 | 0.005798434 |
| PYGL | Glycogen phosphorylase L OS | 1113849.95 | 129708.7053 | 3.373 | 0.001558128 |
| PYCARD | PYD and CARD domain containing OS | 1039041.248 | 99525.0158 | 3.385 | 0.001486423 |
| TOP1 | DNA topoisomerase I OS | 518301.8499 | 168463.4519 | 3.394 | 0.005628887 |
| PTGS1 | Prostaglandin G/H synthase 1 OS | 970772.7778 | 379388.8597 | 3.413 | 0.001200881 |
| F5 | Coagulation factor V OS | 231727.7091 | 5445.31483 | 3.431 | 0.008450255 |
| PPM1K | Protein phosphatase, Mg2+/Mn2+ dependent 1K OS | 172215.5422 | 85457.44587 | 3.449 | 0.006979588 |
| LOC479600 | Calsyntenin 1 OS | 227513.2813 | 65887.87617 | 3.453 | 0.008815083 |
| HCLS1 | Hematopoietic cell-specific Lyn substrate 1 OS | 122314.9434 | 29261.66138 | 3.469 | 0.013604855 |
| MBNL1 | Muscleblind like splicing regulator 1 OS | 1628906.663 | 373233.1585 | 3.472 | 0.000848898 |
| CD34 | Hematopoietic progenitor cell antigen CD34 OS | 1228530.912 | 229226.2077 | 3.485 | 0.001477112 |
| KRT15 | Keratin 15 OS | 5423660.277 | 1685071.048 | 3.492 | 0.001740282 |
| C3 | Complement C3 OS | 49183183.23 | 11487663.3 | 3.513 | 0.002021069 |
| PPID | Peptidylprolyl isomerase D OS | 1807899.512 | 286346.5463 | 3.533 | 0.000705314 |
| RALB | RAS like proto-oncogene B OS | 1541623.23 | 375436.6999 | 3.544 | 0.00145143 |
| CDH5 | Cadherin 5 OS | 77192.41681 | 64669.97422 | 3.547 | 0.014082844 |
| APMAP | Uncharacterized protein OS | 835672.2391 | 120712.6228 | 3.563 | 0.001282882 |
| NAA38 | N-alpha-acetyltransferase 38, NatC auxiliary subunit OS | 1254294.125 | 348801.5646 | 3.596 | 0.001057626 |
| CNTLN | Centlein OS | 532819.8019 | 114442.368 | 3.625 | 0.004303551 |

| | | | | | |
|---|---|---|---|---|---|
| LOC476816 | Uncharacterized protein OS | 4200151.352 | 1027628.288 | 3.625 | 0.001348346 |
| MMP8 | Matrix metallopeptidase 8 OS | 3944832.015 | 177002.4398 | 3.648 | 0.000679325 |
| H1-5 | Uncharacterized protein OS | 212671.3426 | 71167.17747 | 3.657 | 0.005198617 |
| ARPC4 | Actin related protein 2/3 complex subunit 4 OS | 2075851.745 | 217817.1886 | 3.695 | 0.000324506 |
| KRT2 | Keratin, type II cytoskeletal 2 epidermal OS | 476238.3817 | 95562.56025 | 3.74 | 0.007552214 |
| ARPC1B | Uncharacterized protein OS | 1194069.059 | 278861.7272 | 3.762 | 0.000839617 |
| PTPN6 | Protein tyrosine phosphatase non-receptor type 6 OS | 483552.9364 | 17494.40191 | 3.786 | 0.006253909 |
| YWHAQ | Uncharacterized protein OS | 6249128.906 | 1871564.868 | 3.791 | 0.000823082 |
| NPPA | Natriuretic peptides A OS | 33079050.94 | 8871562.201 | 3.834 | 0.000933099 |
| TMEM120A | Transmembrane protein 120A OS | 235564.4436 | 36560.18552 | 3.864 | 0.003213005 |
| STMN1 | Stathmin 1 OS | 1468317.559 | 351980.3707 | 3.873 | 0.000634455 |
| FKBP2 | Uncharacterized protein OS | 1289915.806 | 37114.31836 | 3.878 | 0.000427212 |
| PTPN1 | Protein tyrosine phosphatase non-receptor type 1 OS | 287580.0223 | 64234.21718 | 3.891 | 0.006111366 |
| ABI1 | Uncharacterized protein OS | 1056157.366 | 117396.2979 | 3.897 | 0.000209862 |
| RBX1 | Ring-box 1 OS | 2137651.897 | 512332.735 | 3.909 | 0.000156706 |
| MFAP5 | Microfibril associated protein 5 OS | 440991.0996 | 105102.6655 | 3.918 | 0.006665695 |
| CISD2 | CDGSH iron sulfur domain 2 OS | 2044229.942 | 404143.0431 | 3.935 | 0.00018759 |
| HMGB1 | High mobility group protein B1 OS | 7048223.3 | 3702360.709 | 3.964 | 0.000686899 |
| LOC100855476 | Uncharacterized protein OS | 6969975.853 | 623585.2339 | 4.002 | 0.000501055 |
| A2M | Alpha-2-macroglobulin OS | 3921644.042 | 824135.7097 | 4.028 | 0.000317644 |
| CA2 | Carbonic anhydrase 2 OS | 3580846.087 | 385608.1866 | 4.073 | 0.000221742 |
| LOC119879923 | Alpha-amylase OS | 48348.21629 | 69157.22158 | 4.169 | 0.005931464 |
| ACOT7 | Acyl-CoA thioesterase 7 OS | 1050068.495 | 245050.2223 | 4.194 | 0.000125665 |
| RPL13 | Ribosomal protein L13 OS | 107061.5261 | 71403.70023 | 4.212 | 0.002545293 |
| LCP1 | Lymphocyte cytosolic protein 1 OS | 3372864.715 | 601083.4879 | 4.213 | 0.000139052 |
| APOA1 | Apolipoprotein A-I OS | 311162832.8 | 80289573.9 | 4.254 | 0.00035017 |
| UMPS | Uridine monophosphate synthetase OS | 266970.2922 | 56874.36422 | 4.258 | 0.002624014 |
| IKBIP | IKBKB interacting protein OS | 497294.0297 | 109256.7349 | 4.27 | 0.001773248 |
| CD5L | Apoptosis inhibitor of macrophage OS | 1998715.57 | 236971.091 | 4.299 | 3.31181E-05 |
| GIT2 | GIT ArfGAP 2 OS | 95775.01587 | 22259.24624 | 4.303 | 0.004428792 |
| NES | Nestin OS | 25679294.76 | 5720955.163 | 4.314 | 0.000305586 |
| ALB | Albumin OS | 2534790782 | 764204548.3 | 4.318 | 0.000302913 |
| MT-ATP8 | ATP synthase protein 8 OS | 2782587.239 | 638678.0291 | 4.357 | 4.67208E-05 |
| TIMMDC1 | Translocase of inner mitochondrial membrane domain containing 1 OS | 840888.4232 | 137119.0587 | 4.468 | 0.000102248 |
| SLC12A7 | Solute carrier family 12 member 7 OS | 1256424.955 | 118509.868 | 4.494 | 7.82527E-05 |
| CTSH | Cathepsin H OS | 1961905.334 | 434943.1244 | 4.511 | 2.72058E-05 |
| LOC611458 | Uncharacterized protein OS | 10163323.75 | 953284.2737 | 4.518 | 0.000192637 |
| TTR | Transthyretin OS | 59478323.62 | 12267247.39 | 4.595 | 0.000162304 |
| CTSG | Cathepsin G OS | 19276543.24 | 323246.1131 | 4.62 | 0.000153541 |
| LTBP2 | Latent transforming growth factor beta binding protein 2 OS | 43460.59872 | 9149.820604 | 4.631 | 0.00614464 |
| FGG | Fibrinogen gamma chain OS | 70017266.66 | 8575905.335 | 4.657 | 0.000141456 |
| G6PD | Glucose-6-phosphate dehydrogenase OS | 1243747.709 | 203637.8243 | 4.714 | 7.90348E-05 |
| DNAJC5 | DnaJ heat shock protein family (Hsp40) member C5 OS | 95062.36955 | 20147.47587 | 4.718 | 0.00240711 |
| CEACAM1 | CEA cell adhesion molecule 1 OS | 2367126.25 | 413726.297 | 4.76 | 1.78774E-05 |
| SNX1 | Sorting nexin 1 OS | 399498.9969 | 55464.14942 | 4.768 | 0.001416807 |
| MRPL1 | Uncharacterized protein OS | 1201486.202 | 278576.4974 | 4.812 | 5.92502E-05 |
| MIF | Macrophage migration inhibitory factor OS | 32508278.06 | 4555220.562 | 4.825 | 9.77373E-05 |
| SLC2A3 | Solute carrier family 2, facilitated glucose transporter member 3 OS | 1221299.633 | 43871.32616 | 4.833 | 2.4898E-05 |
| HBQ1 | Uncharacterized protein OS | 1090359.014 | 210251.6959 | 4.855 | 2.24996E-05 |
| C5orf51 | Chromosome 5 open reading frame 51 OS | 901732.9981 | 184842.2019 | 4.878 | 2.64587E-05 |
| PLEKHA7 | Pleckstrin homology domain containing A7 OS | 280718.0014 | 144441.6734 | 4.912 | 0.001102772 |
| MPRIP | Myosin phosphatase Rho interacting protein OS | 5298691.571 | 967514.0664 | 4.958 | 4.38402E-05 |
| STUB1 | STIP1 homology and U-box containing protein 1 OS | 728660.887 | 24147.64771 | 4.997 | 0.000184595 |
| TIMM9 | Translocase of inner mitochondrial membrane 9 OS | 760535.7765 | 150653.4216 | 5.048 | 2.52764E-05 |
| CHCHD1 | Coiled-coil-helix-coiled-coil-helix domain containing 1 OS | 94991.49389 | 6843.70819 | 5.129 | 0.001392116 |
| MRPL48 | Uncharacterized protein OS | 902658.9362 | 142227.1001 | 5.134 | 1.37888E-05 |
| COX17 | Cytochrome c oxidase copper chaperone OS | 653178.7652 | 166328.1124 | 5.147 | 8.39935E-05 |
| FGA | Fibrinogen alpha chain OS | 98594466.17 | 8868616.901 | 5.148 | 4.88077E-05 |
| ADAM10 | ADAM metallopeptidase domain 10 OS | 1912276.045 | 370525.2495 | 5.161 | 4.85912E-06 |
| ASCC3 | Activating signal cointegrator 1 complex subunit 3 OS | 83514.16211 | 16569.15459 | 5.197 | 0.00151309 |
| FGB | Fibrinogen beta chain (Fragment) OS | 4470988.405 | 859411.4485 | 5.202 | 1.77695E-05 |
| AHCYL2 | Adenosylhomocysteinase like 2 OS | 920005.3306 | 175605.9296 | 5.239 | 8.94076E-06 |
| DRG1 | Developmentally regulated GTP binding protein 1 OS | 611410.5516 | 56508.56727 | 5.276 | 0.000184128 |
| PPP1R12C | Protein phosphatase 1 regulatory subunit 12C OS | 811241.2656 | 391610.0062 | 5.329 | 5.07674E-06 |
| MRPL28 | Uncharacterized protein OS | 76668.2165 | 40583.13294 | 5.348 | 0.001109239 |
| ACSL4 | Acyl-CoA synthetase long chain family member 4 OS | 812517.7647 | 126990.3304 | 5.459 | 1.39174E-05 |
| MPO | Myeloperoxidase OS | 22438729.62 | 147715.3627 | 5.473 | 2.47504E-05 |
| PRR32 | Proline rich 32 OS | 1021847.51 | 64856.71337 | 5.518 | 4.7909E-06 |
| TXNDC9 | Thioredoxin domain containing 9 OS | 378406.737 | 150813.9016 | 5.609 | 0.000723519 |

| | | | | | |
|---|---|---|---|---|---|
| WDR48 | WD repeat domain 48 OS | 2403459.125 | 427238.0607 | 5.626 | 1.40955E-06 |
| PRKAG3 | Protein kinase AMP-activated non-catalytic subunit gamma 3 OS | 2694904 | 476781.4935 | 5.652 | 1.04698E-06 |
| ITGB2 | Integrin subunit beta 2 OS | 2625761.742 | 141828.288 | 5.798 | 9.7214E-07 |
| LYN | LYN proto-oncogene, Src family tyrosine kinase OS | 956172.3492 | 164513.4405 | 5.812 | 1.97773E-06 |
| HBQ1 | Uncharacterized protein OS | 1240944732 | 246399991.5 | 5.852 | 1.14366E-05 |
| CD177 | Uncharacterized protein OS | 9077013.382 | 44976.29632 | 5.853 | 1.28341E-05 |
| CCN3 | Cellular communication network factor 3 OS | 1253050.069 | 55378.45258 | 5.894 | 1.55516E-06 |
| LOC100855 538 | Ribosomal protein S24 OS | 162668.3245 | 27342.86791 | 5.949 | 9.55544E-05 |
| MRPS31 | Mitochondrial ribosomal protein S31 OS | 1699982.777 | 275434.342 | 6.166 | 1.46079E-06 |
| FGB | Fibrinogen beta chain OS | 68465841.54 | 9792468.997 | 6.284 | 4.88881E-06 |
| FHL1 | Uncharacterized protein OS | 51080482.79 | 11652216.2 | 6.297 | 4.76495E-06 |
| RPL36AL | Ribosomal protein L36a like OS | 1247656.965 | 116284.908 | 6.321 | 6.17898E-07 |
| LOC480784 | Uncharacterized protein OS | 603946.068 | 104846.0438 | 6.346 | 1.47868E-05 |
| CORO1A | Coronin 1A OS | 2804540.255 | 217793.8449 | 6.436 | 2.25136E-07 |
| Lysozyme | Lysozyme C, spleen isozyme OS | 4796196.529 | 122335.4752 | 6.46 | 6.81947E-07 |
| NCF2 | Neutrophil cytosolic factor 2 OS | 1452178.953 | 210143.7362 | 6.462 | 8.51105E-07 |
| CRIP1 | Uncharacterized protein OS | 65475642.59 | 9104305.199 | 6.495 | 3.25917E-06 |
| LOC119863 903 | Uncharacterized protein OS | 20492213.63 | 3407499.501 | 6.621 | 2.56928E-06 |
| PSTPIP1 | Proline-serine-threonine phosphatase interacting protein 1 OS | 659823.9346 | 52845.84818 | 6.725 | 7.31848E-06 |
| STAM | Signal transducing adaptor molecule OS | 793041.375 | 63804.09022 | 6.824 | 4.39101E-06 |
| EMID1 | EWS RNA binding protein 1 OS | 332131.6796 | 38582.12985 | 7.145 | 5.62808E-05 |
| HBQ1 | Hemoglobin subunit alpha1 OS | 9380359.514 | 792166.952 | 7.149 | 1.11952E-06 |
| MANF | Mesencephalic astrocyte derived neurotrophic factor OS | 1269665.686 | 268929.8849 | 7.53 | 1.02862E-07 |
| LTF | Lactotransferrin OS | 26546171.35 | 137722.5356 | 7.665 | 3.86143E-07 |
| DDX39A | DExD-box helicase 39A OS | 577854.625 | 71533.93512 | 8.078 | 1.78468E-06 |
| BPI | Bactericidal permeability increasing protein OS | 2572636.528 | 106717.6551 | 8.495 | 1.4799E-09 |
| BCAP31 | B cell receptor associated protein 31 OS | 1642516.404 | 371004.5606 | 8.496 | 5.22295E-09 |
| GRK6 | G protein-coupled receptor kinase 6 OS | 401369.7344 | 46801.32779 | 8.576 | 1.52098E-05 |
| ELANE | Elastase, neutrophil expressed OS | 16754454.16 | 531466.5875 | 8.609 | 7.79671E-08 |
| LIMS2 | LIM zinc finger domain containing 2 OS | 156168.6166 | 25985.14078 | 8.718 | 3.45845E-06 |
| BOLA3 | Uncharacterized protein OS | 964549.3809 | 100132.7388 | 8.758 | 3.05922E-09 |
| HBB | Hemoglobin subunit beta OS | 335763471.6 | 40235080.39 | 8.934 | 4.59401E-08 |
| SERBP1 | Uncharacterized protein OS | 714119.7717 | 54444.30183 | 9.006 | 1.42531E-07 |
| KCNJ3 | Potassium inwardly rectifying channel subfamily J member 3 OS | 326098.1563 | 80585.9317 | 9.27 | 4.12407E-06 |
| ACOT6 | Acyl-CoA thioesterase 6 OS | 6110658.135 | 492228.1406 | 9.666 | 5.66059E-09 |
| NFIC | Nuclear factor I C OS | 305904.1868 | 28254.00687 | 10.827 | 4.27495E-07 |
| NEBL | Nebulette OS | 517859.2604 | 47781.60942 | 10.838 | 7.65912E-07 |
| ARPC5 | Actin related protein 2/3 complex subunit 5 OS | 5851704 | 525341.7758 | 11.139 | 8.7272E-10 |
| Cytochrome | Cytochrome c oxidase subunit 6C OS | 7073558.802 | 951280.896 | 11.23 | 7.63549E-10 |
| ABLIM1 | Actin binding LIM protein 1 OS | 303337.3638 | 26092.11331 | 11.626 | 3.15369E-07 |
| TCEAL4 | Uncharacterized protein OS | 327018.315 | 25445.50903 | 12.516 | 1.81762E-07 |
| GABARAPL 2 | GABA type A receptor associated protein like 2 OS | 572880.0803 | 24270.19828 | 13.874 | 1.54881E-08 |
| DDX46 | DEAD-box helicase 46 OS | 169691.8907 | 12422.48113 | 14.261 | 1.25769E-08 |
| S100A8 | Protein S100 OS | 18884957.66 | 730255.1724 | 15.242 | 8.61156E-12 |
| TPM3 | Tropomyosin 3 OS | 2131037.002 | 67127.97524 | 15.39 | 7.77156E-15 |
| CAMP | Cathelicidin OS | 31155873.49 | 631262.9411 | 18.115 | 3.64153E-13 |
| PPP3CC | Protein phosphatase 3 catalytic subunit gamma OS | 8298920.495 | 685420.5608 | 21.014 | 3.84137E-14 |
| RAP1B | RAP1B, member of RAS oncogene family OS | 2168958.787 | 58958.0935 | 21.161 | 1E-17 |
| DNAJC19 | DnaJ heat shock protein family (Hsp40) member C19 OS | 4036339.869 | 184709.1746 | 21.852 | 2.22045E-16 |
| THBS4 | Thrombospondin 4 OS | 733015.8596 | 28292.13541 | 25.909 | 4.44089E-15 |
| MMGT1 | Uncharacterized protein OS | 5027189.302 | 187525.3564 | 26.808 | 1E-17 |
| ME1 | Malic enzyme 1 OS | 34550383.48 | 601888.2037 | 42.235 | 1E-17 |
| CRELD1 | Cysteine rich with EGF like domains 1 OS | 747506.905 | 17433.84821 | 42.877 | 1E-17 |
| MYOM1 | Myomesin 1 OS | 875431.8167 | 13620.05519 | 64.275 | 1E-17 |
| GGH | Gamma-glutamyl hydrolase OS | 820066.3438 | 2363.882009 | 91.096 | 1E-17 |
| ACOX3 | Acyl-CoA oxidase 3, pristanoyl OS | 350012.9974 | 0 | 100 | 1E-17 |
| ACTR2 | Actin related protein 2 OS | 643147.2046 | 0 | 100 | 1E-17 |
| ALAS1 | 5'-aminolevulinate synthase 1 OS | 541697.4743 | 0 | 100 | 1E-17 |
| ALDH3A1 | Aldehyde dehydrogenase, dimeric NADP-preferring OS | 790265.7465 | 0 | 100 | 1E-17 |
| ARIH2 | Ariadne RBR E3 ubiquitin protein ligase 2 OS | 89230.38851 | 0 | 100 | 1E-17 |
| ROCK1 | Rho associated coiled-coil containing protein kinase 1 OS | 122674.7042 | 0 | 100 | 1E-17 |
| CIAO3 | Cytosolic iron-sulfur assembly component 3 OS | 52259.66038 | 0 | 100 | 1E-17 |
| CPE | Carboxypeptidase E OS | 307606.1965 | 0 | 100 | 1E-17 |
| CPT1A | Carnitine palmitoyltransferase 1A OS | 831520 | 0 | 100 | 1E-17 |
| FDXR | Ferredoxin reductase OS | 480993.7645 | 0 | 100 | 1E-17 |
| GFAP | Glial fibrillary acidic protein OS | 681011.8125 | 0 | 100 | 1E-17 |
| GRIPAP1 | GRIP1 associated protein 1 OS | 347873.44 | 0 | 100 | 1E-17 |

| | | | | | |
|---|---|---|---|---|---|
| Gypc | Glycophorin C OS | 1554823.218 | 0 | 100 | 1E-17 |
| HCK | HCK proto-oncogene, Src family tyrosine kinase OS | 302837.7319 | 0 | 100 | 1E-17 |
| HNRNPLL | Heterogeneous nuclear ribonucleoprotein L like OS | 309406.6893 | 0 | 100 | 1E-17 |
| HOGA1 | 4-hydroxy-2-oxoglutarate aldolase 1 OS | 1219062.239 | 0 | 100 | 1E-17 |
| KIF1B | Uncharacterized protein OS | 536876.8946 | 0 | 100 | 1E-17 |
| KPNA3 | Karyopherin subunit alpha 3 OS | 238612.5517 | 0 | 100 | 1E-17 |
| KRT15 | Keratin 15 OS | 1113587.974 | 0 | 100 | 1E-17 |
| LDLR | Low density lipoprotein receptor OS | 587342.4283 | 0 | 100 | 1E-17 |
| LSP1 | Lymphocyte specific protein 1 OS | 1789517 | 0 | 100 | 1E-17 |
| MAP2K4 | Mitogen-activated protein kinase kinase 4 OS | 1673056.402 | 0 | 100 | 1E-17 |
| MSRB2 | Methionine sulfoxide reductase B2 OS | 258619.9719 | 0 | 100 | 1E-17 |
| MTMR7 | Myotubularin related protein 7 OS | 203941.5606 | 0 | 100 | 1E-17 |
| NDUFA2 | Uncharacterized protein OS | 1055776.039 | 0 | 100 | 1E-17 |
| NEDD4L | NEDD4 like E3 ubiquitin protein ligase OS | 539941.5037 | 0 | 100 | 1E-17 |
| NME1 | Nucleoside diphosphate kinase A OS | 322696.3125 | 0 | 100 | 1E-17 |
| NUP43 | Nucleoporin 43 OS | 397195.9587 | 0 | 100 | 1E-17 |
| PEAK3 | PEAK family member 3 OS | 353437.1686 | 0 | 100 | 1E-17 |
| PEX26 | Tubulin alpha 8 OS | 1026218.644 | 0 | 100 | 1E-17 |
| PPP1CA | Serine/threonine-protein phosphatase PP1-alpha catalytic subunit OS | 3229891.587 | 0 | 100 | 1E-17 |
| PRRC2A | Proline rich coiled-coil 2A OS | 762167.8736 | 0 | 100 | 1E-17 |
| PYROXD2 | Pyridine nucleotide-disulphide oxidoreductase domain 2 OS | 595097.265 | 0 | 100 | 1E-17 |
| RPS25 | Ribosomal protein S25 OS | 918277.5933 | 0 | 100 | 1E-17 |
| SBSPON | Somatomedin B and thrombospondin type 1 domain containing OS | 431445.6493 | 0 | 100 | 1E-17 |
| SCAMP2 | Secretory carrier membrane protein 2 OS | 195257.1663 | 0 | 100 | 1E-17 |
| SDHAF1 | Succinate dehydrogenase complex assembly factor 1 OS | 768575.5684 | 0 | 100 | 1E-17 |
| SEPTIN6 | Septin 6 OS | 1937786.296 | 0 | 100 | 1E-17 |
| SRSF5 | Serine and arginine rich splicing factor 5 OS | 139734.7077 | 0 | 100 | 1E-17 |
| SUMF2 | Sulfatase modifying factor 2 OS | 218615.2316 | 0 | 100 | 1E-17 |
| TCEA1 | Transcription elongation factor A1 OS | 2130567.816 | 0 | 100 | 1E-17 |
| TCN1 | Uncharacterized protein OS | 496776.4063 | 0 | 100 | 1E-17 |
| UPP1 | Uncharacterized protein OS | 550913.1323 | 0 | 100 | 1E-17 |
| ANXA6 | Annexin A6 OS | 235503.2282 | 0 | 100 | 1E-17 |

***Supplemental Table 2:*** *Significantly upregulated genes in Atrial Fibrillation*

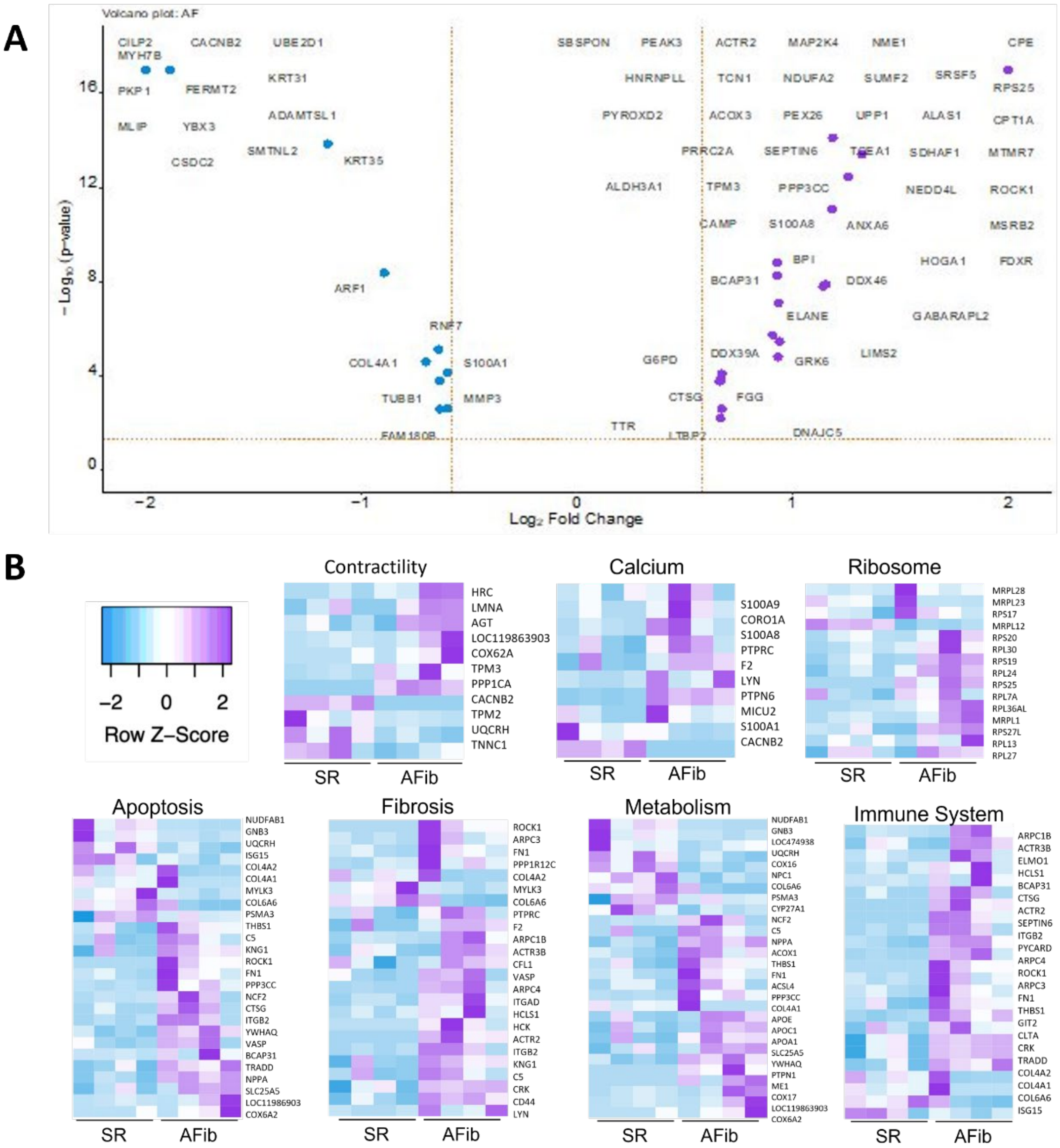


***Supplemental Figure 1:** A)* Volcano Plot of all dysregulated proteins in AFib. B) Other dysregulated Gene Ontology pathways in Atrial Fibrillation canine model and the genes involved in those pathways.

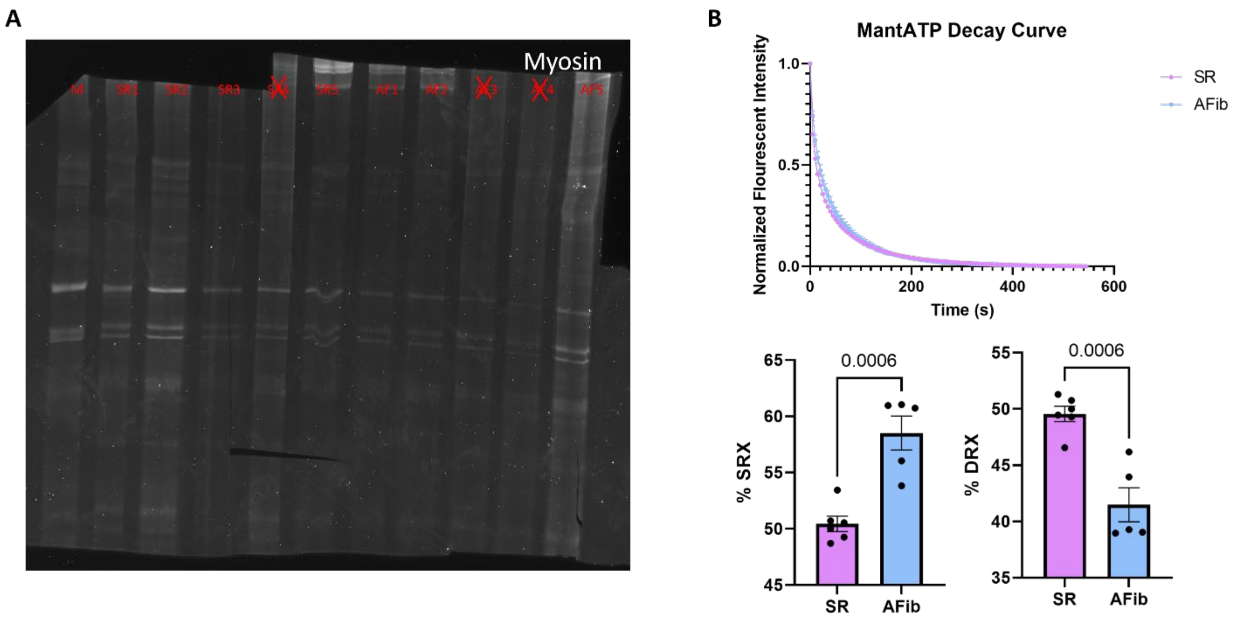


***Supplemental Figure 2:*** *A) Raw image of the myosin gel. B) MantATP fluorescent decay fit to a double exponential decay curve of the SR and AFib tissue shows alterations in both the fast and slow phase of fluorescent decay. Analysis of this fluorescent decay shows that there is an increase in the percent of myosin heads in the SRX conformation and a decrease in the percentin the DRX conformation in the AFib tissue.*

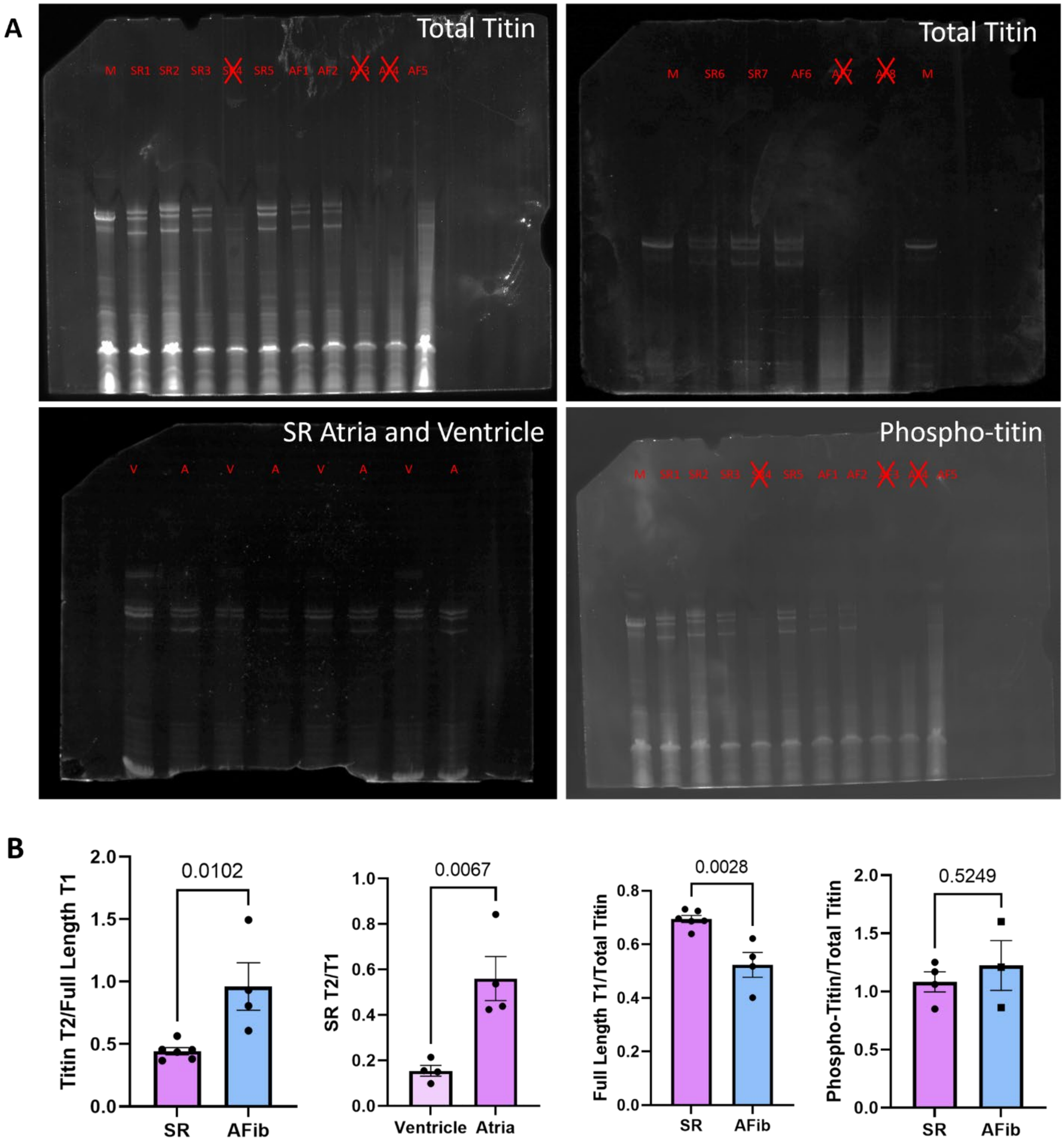


***Supplemental Figure 3:*** *A) Raw images of all titin gels. B) Analysis of titin gels shows a significant increase in T2/T1 ratio and SR ventricle T2/T1 ration, and a significant decrease in the full length T1/Total Titin ration. Analysis of titin phosphorylation shows no difference in total titin phosphorylation between the SR and AFib samples.*

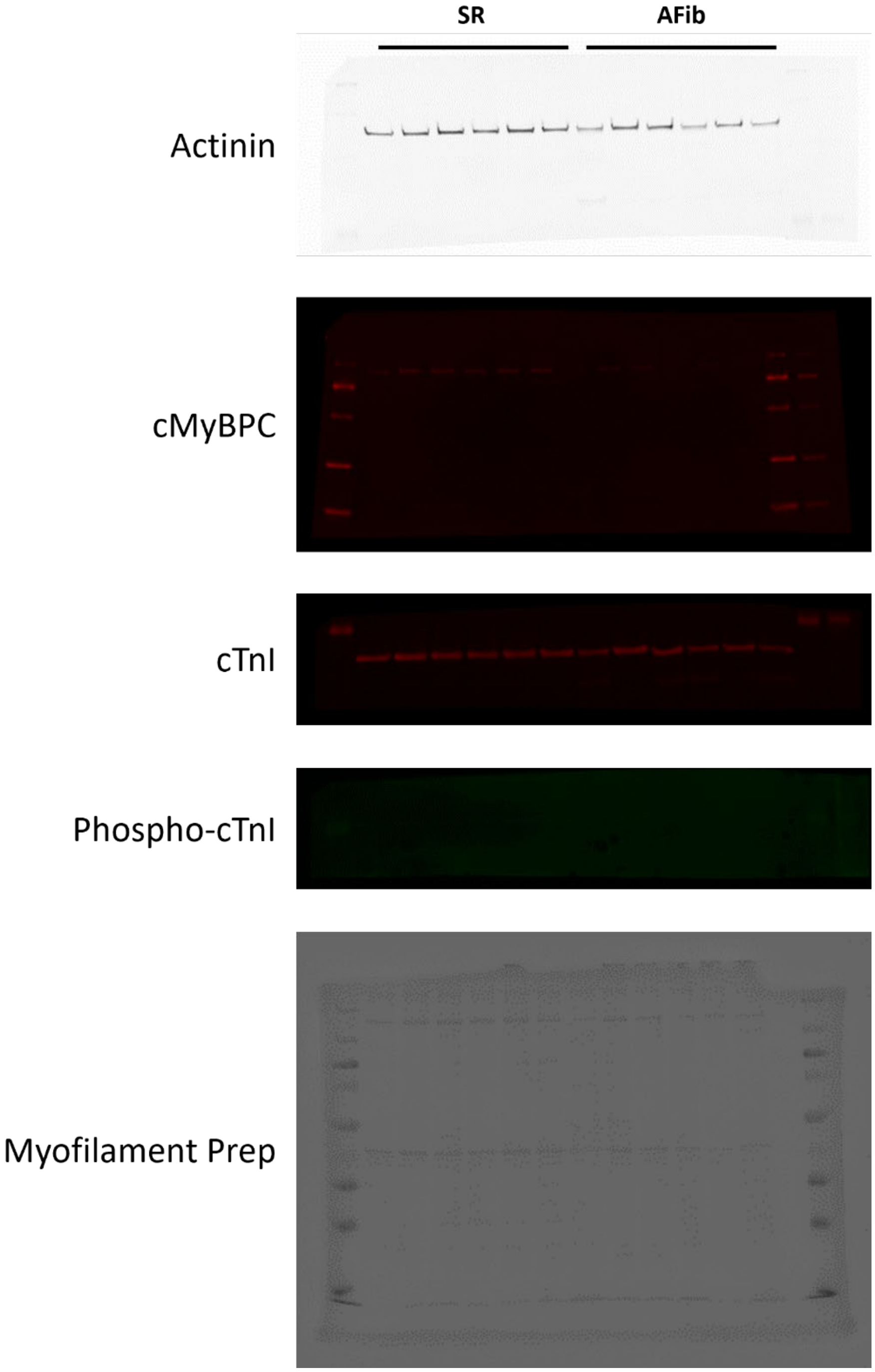


***Supplemental Figure 4:*** All raw images of the immunoblots

**SUPPLEMENTARY METHODS**

*Mass Spectrometry and Data Analysis*. Mass spectrometry and analysis was done as described previously with some modifications (16). Purified peptides from eight canines (n = 4 SR,4 AFib), 1.0 µg, were loaded onto a Vanquish Neo UHPLC system (Thermo Fisher) with a heated trap and elute workflow with a c18 PrepMap, 5 mm, 5 µM trap column (Thermo Fisher #160454) in a forward-flush configuration connected to a 25 cm Easyspray analytical column (Thermo Fisher #ES802A rev2) 2 µM, 100 A, 75 µm x 25 with 100% Buffer A (0.1% Formic acid in water) and the column oven operating at 35°C. Peptides were eluted over a 150 min gradient, using 80% acetonitrile, 0.1% formic acid (buffer B), going from 4% to 15% over 30 min, to 40% over 90 min, then to 65% over 20 min, and then kept at 95% for 10 min, after which all peptides were eluted. Spectra were acquired with an Orbitrap Eclipse Tribrid mass spectrometer with FAIMS Pro interface (Thermo Fisher) running Tune 3.5 and Xcalibur 4.5. For all acquisition methods, spray voltage set to 1900 V, and ion transfer tube temperature set at 300°C, FAIMS switched between CVs of −45 V, – 55 V, and −65 V with cycle times of 1 s. MS1 spectra were acquired at 120,000 resolutions with a scan range from 375 to 1600 m/z, normalized AGC target of 100%, and maximum injection time of 50 ms, S-lens RF level set to 30, without source fragmentation and datatype positive and profile; Precursors were filtered using monoisotopic peak determination set to peptide; included charge states, 2 – 7 (reject unassigned); dynamic exclusion enabled, with n = 1 for 60 s exclusion duration at 10 ppm for high and low. DDMS2 scan using isolation mode Quadrupole, Isolation Window (m/z): 1.6; Activation Type set to HCD with 30% Collision Energy (CE), Detector Type: Ion Trap; Scan Rate: Turbo; AGC Target: 10000; Maximum Injection Time:35 ms, Microscans: 1 and Data Type: Centroid.

*Protein isolation and gels for Titin and Myosin.* Proteins were isolated from all 12 of the canine atria samples (n = 6 SR, 6 AFib) as described before (18). 10 – 15 mg of the frozen tissue was measured from each heart and crushed while maintained in liquid nitrogen. Once fully crushed, samples were incubated at -20 (degrees C) for 20 minutes to slowly bring to back to room temperature. A solution of 8 M urea (8 M Urea, 1 M Thiourea, 3% SDS, 0.05 M Tris HCL, pH 6.8) was added to the ground samples to dissolve them and then 50% glycerol with protease and phosphatase inhibitors was added to preserve protein integrity and stabilize the sarcomeres. The solution was incubated at 60°C to ensure that the tissue is fully dissolved, centrifuged at 21.1k x g for 5 minutes to remove debris from the solution, and the purified supernatant containing the myofilament proteins was collected and flash frozen. Protein concentration was determined using the RC/DC protein quantification assay following manufacturer protocol (Biorad, 5000122) and samples were diluted to 0.5 mg/ml.

Titin isoform expression was evaluated as described before (18). To separate titin isoforms, a 16 cm by 18 cm 1% agarose (Acrylamide Plug: 4 ml 30% 37.5:1 Acrylamide, 2 ml 5X Running Buffer (0.25 M Tris Base, 1.95 M Glycine, 0.05 W/V 10% SDS), 3.89 ddH2O, 100 ul 10% APS, 8.75 ul TEMED; Agarose Gel: 15 ml Glycerol, 10 ml 5x Running Buffer, 0.5 g SeaKem Gold Agarose (Lonza #50150), 24 ml dd H2O) gel was made. 10 µg of each sample in 2x loading dye (n = 6 SR, 4 AFib) was loaded into the gel. The gels were run at 15 mA (one gel) or 30 mA (two gels) for 3 hours and 20 minutes. The gels were then fixed in a 20% methanol, 10% acetic acid solution for one hour, then stained with Sypro Ruby for total titin or ProQ Diamond for phosphorylated titin overnight. The gels were destained for 30 minutes in a 20% methanol solution and imaged using UV imaging (C-600 Azure).

Myosin isoforms were separated, as described before (19), using a Hoefer-format SDS-PAGE gel at 6.25% acrylamide (16.55 ml ddH2O, 300 ul 10% SDS, 5.65 ml 2M Tris pH 8.8, 7.5 ml 25% Acrylamide 99:1 (reagents combined and then degassed for 20 minutes), 312 ul 10% APS W/V, 12.5 ul TEMED). 10 µg of the myofilament protein solution was mixed with 2x loading dye and loaded into the acrylamide gel (n = 4 SR, 3 AFib). The gel was run at 12.5 mA for 20 minutes followed by 15 mA for 21 hours (one gel) or 25 mA for 20 minutes followed by 30 mA (two gels). The gels were then fixed in a 7% acetic acid and 40 % methanol solution, stained with Syro Ruby overnight, and destained in the 7% acetic acid and 40% methanol solution. Once destained, the gels were imaged using UV imaging (Azure c-600).

*Myofilament fractionation and immunoblots.* Tissue samples from all 12 canines (n = 6 SR, 6 Afib) were enriched for myofilament proteins via myofilament fractionation. For myofilament fractionation, tissue was homogenized in F-60 buffer (60 mM KCl, 30 mM Imidazole, 2 mM $MgCL_2$) + protease inhibitor and then spun down at 12,000 g for 10 minutes and the supernatant was discarded. This removed the soluble protein fraction, leaving the myofilament proteins in the pellet. The pellet was washed with F-60 buffer + protease inhibitor + 0.1% triton X-100 3 times. The pellet was then homogenized in RIPA + protease inhibitor to solubilize the myofilament proteins and diluted to 0.2 mg/mL in 2x loading dye.

Total protein or myofilament samples were then run on immunoblots to probe for proteins. Briefly, 10% acrylamide gels were made (Resolving Gel: 1.9 ml H2O, 1.3 ml 1.5 Mtris-Cl pH 8.8, 1.7 ml 30% 37.5:1 Acrylamide, 50 ul 10% SDS, 50 ul 10% APS, 2 ul TEMED; Stacking Gel: 680 ul H2O, 130 ul 1 M Tris-Cl pH 6.8, 170 ul 30% 37.5:1 Acrylamide, 10 ul 10% SDS, 10 ul 10% APS, 1 ul TEMED) and loaded with 7 µg of the chameleon duo pre-stained protein ladder (Licor, 928-60000) and 12 µg of each sample (n = 6 SR, 6 AFib). This was run at 100 V for 1.5 hours and then transferred to a membrane at 300 mA for 3 hours. The membrane was blocked in 1% casein blocker (Bio-Rad, 1610782) then soaked overnight in primary antibodies: TnI 1:500 (Santa Cruz, 365446); P-TnI 1:500 (Cell Signaling, 4004S); cMyBPC 1:2500 (Santa Cruz, 137180); MHC 1:2500 (Developmental Studies Hybridoma Bank, MF20); ACTN2 1:5000 (Proteintech, 14221-1-AP). The membrane was washed using TBS-T, stained with either Goat anti-Mouse 680 1:5000 (Azure AC2129) or Goat anti-Rabbit 800 1:5000 (Azure AC2134) secondary antibodies, and imaged using near infra-red imaging (Azure c-600). Once imaged, the blot was stained for total protein using reversible stain and dried out. All protein gels were analyzed via densitometry in ImageJ version 2.9 and significance was determined using T-Tests

*MantATP Assay.* Canine cardiac ventricle tissue (5 mg) was flash frozen in liquid nitrogen and broken into smaller pieces. MantATP protocol was adapted from a previous publication (*add McNamara* citation). The tissue pieces were permeabilized and rocked in Skinning buffer (in mM) (100 NaCl, 8 $MgCl_2$, 5 EGTA, 5 $K_2PO_4$, 5 $KH_2PO_4$, 3 $NaN_3$, 5 ATP, 1 DTT, 20 BDM, and 0.01% Triton X-100 at pH 7) with 2-hour washes for 6 hours total at 4°C. After 6 hours, Skinning buffer was changed to Glycerinating buffer (in mM) (120 K Acetate, 5 Mg Acetate, 5 EGTA, 2.5 $K_2PO_4$, 2.5 $KH_2PO_4$, 50 MOPS, 5 ATP, 20 BDM, 2 DTT, 50% Glycerol at pH 6.8) and continued to rock overnight at 4°C, and then refreshed with glycerinating buffer in the morning.

Tissue was pulled to size in cold glycerinating buffer and secured onto a glass coverslip. Tissue was washed with Rigor buffer 1 (120 K Acetate, 5 Mg Acetate, 5 EGTA, 2.5 $K_2PO_4$, 2.5 $KH_2PO_4$, 50 MOPS, and 2 fresh DTT at pH 6.8) 10 minutes before imaging. Rigor buffer 1 was then exchanged with Rigor buffer 2 (Rigor buffer 1+ 0.25 mM mantATP) and images were taken every 5 seconds at 72 ms exposure for 9 minutes using a Nikon Eclipse T*i* scope with 20X objective. After 9 minutes of imaging, Relaxing solution (Rigor buffer 1+ 4 mM ATP) was added onto the canine tissue and imaging continued for another 9 minutes.

ImageJ was used to measure fluorescence intensity in 6 different regions of the canine tissue and data was fit using a double exponential decay formula where *I* is fluorescence intensity.

$$I = 1 - P_1\left(1 - \frac{-t}{e^{T_1}}\right) - P_2\left(1 - \frac{-t}{e^{T_2}}\right)$$

A non-specific binding factor was calculated and subtracted from initial fluorescence to find $P_2$ (SRX). Statistical analysis was then performed using a T-Test, along with curve fitting, in GraphPad PRISM v. 9.
